# The Aspartic Protease Ddi1 Contributes to DNA-Protein Crosslink Repair in Yeast

**DOI:** 10.1101/575860

**Authors:** Nataliia Serbyn, Audrey Noireterre, Ivona Bagdiul, Michael Plank, Agnès H Michel, Robbie Loewith, Benoît Kornmann, Françoise Stutz

**Author notes:** Correspondence (F.S.), (N.S.). Department of Biochemistry, University of Oxford, Oxford OX1 3QU, United Kingdom. These authors contributed equally (Michael Plank, Agnès H Michel).

## Abstract

Naturally occurring or drug-induced DNA-protein crosslinks (DPCs) interfere with key DNA transactions if not timely repaired. The unique family of DPC-specific proteases Wss1/SPRTN targets DPC protein moieties for degradation, including topoisomerase-1 trapped in covalent crosslinks (Top1ccs). Here we describe that the efficient DPC disassembly requires Ddi1, another conserved predicted protease in *Saccharomyces cerevisiae*. We found Ddi1 in a genetic screen of the *tdp1wss1* mutant defective in Top1cc processing. Ddi1 is recruited to a persistent Top1cc-like DPC lesion in an S-phase dependent manner to assist eviction of crosslinked protein from DNA. Loss of Ddi1 or its putative protease activity hypersensitize cells to DPC trapping agents independently from Wss1 and 26S proteasome, implying its broader role in DPC repair. Among potential Ddi1 targets we found the core component of RNAP II and show that its genotoxin-induced degradation is impaired in *ddi1.* Together, we propose that the Ddi1 protease contributes to DPC proteolysis.

## INTRODUCTION

Genuine cell division requires efficient duplication of both DNA strands, which in turn demands the unimpeded progression of the replication fork. DNA lesions or physical barriers such as DNA-protein crosslinks (DPCs) drastically slow the progression of eukaryotic replicative CMG helicase, consequently inhibiting DNA synthesis (Stingele et al., 2017). Diverse endogenous, environmental and therapeutic agents can irreversibly covalently trap proteins in DPCs. The crosslinking agents include but are not limited to reactive aldehydes, ionizing radiation, UV light, or metal ions (Ide et al., 2018). Several spontaneous or drug-induced abortive enzymatic reactions catalyzed by topoisomerases, PARP1, DNA Polβ, DNMT1, etc, can also result in the formation of DPCs (Klages-Mundt and Li, 2017; Stingele et al., 2017). Canonical repair pathways, including homologous recombination (HR) and nucleotide excision repair (NER), are required for cell survival in the presence of DPCs (de Graaf et al., 2009; Nakano et al., 2009). However, the access of repair factors to the lesions could be limited by the bulkiness of the DPC; for example, NER in mammals only removes crosslinked peptides smaller than 8-10kDa (Nakano et al., 2009). DNA proximal to the adduct can be further cleaved by the MRN complex (Deshpande et al., 2016; Hoa et al., 2016) or by structure-specific endonucleases such as budding yeast Rad1-Rad10, Slx1-Slx4 or Mus81-Mms4 complexes (Pommier et al., 2006). In both cases, prior size reduction or complete elimination of the adduct’s protein moiety would relieve the steric hindrance, thus facilitating access to the damage site (Ide et al., 2018).

Eukaryotic cells carry at least one dedicated DPC protease that diminishes the bulkiness of DPCs. Recent discoveries shed light on its molecular identity, describing the metalloprotease Wss1 in *Saccharomyces cerevisiae* (Balakirev et al., 2015; Stingele et al., 2014) and its homologue in higher eukaryotes SPRTN, also known as DVC1 (Lopez-Mosqueda et al., 2016; Maskey et al., 2017; Stingele et al., 2016; Vaz et al., 2016), as the DPC-specific proteases. Both Wss1 and SPRTN are activated by DNA and cooperate with the Cdc48/p97 segregase to proteolyze DPCs (Stingele et al., 2015). Despite similar properties, Wss1 and SPRTN could be distinctly regulated in yeast and mammals, as confirmed by the observation that SPRTN cannot substitute yeast Wss1 in Top1cc repair (Lopez-Mosqueda et al., 2016). Wss1 has a SUMO-interacting motif and is proposed to be regulated by HMW-SUMO species (Balakirev et al., 2015; Stingele et al., 2014). SPRTN, in contrast, possesses ubiquitin-binding motifs, binds PCNA, is cell-cycle regulated and activated by de-ubiquitination and ssDNA (Larsen et al., 2018; Mosbech et al., 2012; Stingele et al., 2016). In addition to the dedicated proteolytic mechanism, it was proposed previously and confirmed recently that the 26S proteasome is directly involved in DPC proteolysis (Klages-Mundt and Li, 2017; Larsen et al., 2018; Lin et al., 2008; Quievryn and Zhitkovich, 2000). In addition, the CMG helicase can bypass DPCs with the help of RTEL1 prior to proteolysis, however SPRTN and 26S proteasome activities remain the prerequisite for the full performance of a downstream repair pathway (Sparks et al., 2019). In this case SPRTN-mediated proteolysis occurs behind the replication fork in post-replicative manner and is independent of DPC ubiquitination (Larsen et al., 2018).

One class of proteins prone to DPC formation comprises DNA topoisomerases, highly conserved enzymes essential to relax torsional stress (Ide et al., 2018; Klages-Mundt and Li, 2017; Stingele et al., 2017). Topoisomerase 1 (Top1) nicks one DNA strand and forms a transient covalent Top1-DNA intermediate allowing DNA rotation (Pommier et al., 2006). Whereas these transient ssDNA breaks are usually efficiently rejoined, impaired ligation can trap Top1 on 3’ DNA thus stabilizing a covalent crosslink (cc) – Top1cc. Formation of Top1ccs can be induced by camptothecin (CPT) and its derivatives, the drugs extensively used in anticancer therapy, which prevent DNA re-ligation by entering Top1 catalytic pocket (Pommier et al., 2006). Crosslinked Top1 can be directly extracted from DNA by Tdp1 phosphodiesterase (Connelly and Leach, 2004). In the absence of Tdp1 that hydrolyzes the Top1-DNA phosphotyrosyl link, Top1cc proteolysis by Wss1 is nearly essential for *S. cerevisiae* survival. This observation allowed the initial classification of Wss1 as a DPC protease (Stingele et al., 2014). Thus, Top1ccs proved to be a good model to discover novel DPC metabolism factors.

In this study we identified *DDI1* in a genetic screen using the *tdp1wss1* mutant defective for Top1cc processing. Initially discovered in yeast as a gene overexpressed from a bidirectional promoter in response to methyl methanesulfonate treatment (Liu and Xiao, 1997), <u>D</u>NA-<u>D</u>amage Inducible-<u>1</u> (Ddi1) was further proposed to function in numerous processes, including v-SNARE binding, regulation of protein secretion, mitotic checkpoint control and mating type switch (Clarke et al., 2001; Kaplun et al., 2005; Lustgarten and Gerst, 1999). Mammalian cells have two homologues of the yeast gene, *DDI1* and *DDI2*, that were previously shown to control proteasomal gene expression (Koizumi et al., 2016) and to remove RTF2 from HU-stalled replication forks (Kottemann et al., 2017). In *S. cerevisiae*, Ddi1 together with Rad23 and Dsk2 belongs to the ubiquitin receptor family comprising the scaffolds that mediate interaction between the proteasome and its ubiquitinated substrates (Dantuma et al., 2009). The N-terminal ubiquitin-like (UBL) and the C-terminal ubiquitin associated (UBA) domains of Ddi1, the classical signature of this family, account for most of the Ddi1 functions mentioned above. Nevertheless, the UBL domain has a weak sequence identity with other members of the ubiquitin receptor family (Nowicka et al., 2015); furthermore, the UBA domain of yeast Ddi1 is not evolutionary conserved and is completely lost in the vertebrate branch (Siva et al., 2016). On the other hand, the central part of Ddi1 encodes a highly conserved retroviral protease (RVP) domain with an HIV-like aspartic protease fold, emphasizing that Ddi1 is rather an unusual ubiquitin receptor (Siva et al., 2016; Trempe et al., 2016). Consistent with the conservation of the RVP domain among eukaryotes, structural studies demonstrated that the Ddi1 dimer adopts a typical aspartic protease fold, strongly suggesting that the yeast and mammalian enzymes must be proteolytically active (Sirkis et al., 2006; Siva et al., 2016; Trempe et al., 2016). Although the Ddi1 proteolytic activity was previously addressed, it was not yet confirmed *in vitro* (Siva et al., 2016; Trempe et al., 2016). The functional relevance of Ddi1 protease activity therefore remains elusive.

Here we report that the putative protease activity of yeast Ddi1 is required for tolerance towards DNA-protein crosslinks, including Top1 poisons. We show that Ddi1 facilitates the removal of Top1cc-mimicking DPCs induced by a mutant Flp recombinase. Ddi1 functions redundantly with Wss1 to resolve a variety of enzymatic and chemical-induced DPCs and helps to remove stalled RNA polymerase II (RNAP II) from chromatin in a protease-dependent manner. Together, we propose that Ddi1 is a novel protease that actively participates in DPC repair in budding yeast.

## RESULTS

### The Ddi1 aspartic protease provides resistance towards Top1-DNA crosslinks

Aiming to discover additional Top1-DNA crosslink repair factors, we designed an unbiased genetic screen of a budding yeast mutant impaired in Top1cc processing. Co-depletion of the Tdp1 phosphodiesterase and the Wss1 metalloprotease in *S. cerevisiae*, causes a growth defect due to Top1cc accumulation (Balakirev et al., 2015; Stingele et al., 2014). The severely impaired *tdp1*Δ*wss1*Δ double mutant rapidly accumulates suppressor mutations. To overcome this limitation, we transiently depleted Tdp1 protein using the auxin-inducible degron (AID) system (Morawska and Ulrich, 2013). When fused to a short version of the AID* tag, Tdp1 was efficiently degraded in the presence of auxin (Figures 1A and S1A). The Tdp1 degron (*Tdp1-AID*-6HA pADH1-TIR1*) in combination with *wss1*Δ negatively impacted the growth of the strain and sensitized it to the Top1 trapping drug CPT, similar to loss of both proteins (Figure S1B). The Tdp1 degron combined with *wss1*Δ (for the simplicity hereafter referred to as *tdp1wss1*) was further used to generate a SATAY library of random transpositions (Michel et al., 2017) in the absence of auxin, followed by Tdp1 depletion and mapping of the transposition events by sequencing (Figure 1B). To identify synthetic rescue and lethality *tdp1wss1* genetic interactions, we compared the number of transposons mapping in each of the 6600 yeast ORFs in this library against a pool of six previously-published SATAY libraries (Michel et al., 2017) and represented the results as a Volcano plot (Figure 1C). As expected, the *TOP1* gene was a key suppressor (Figure 1C), confirming the robustness of the screen and validating previous observations, which described Top1 poisons as the principle source of *tdp1*Δ*wss1*Δ growth defect (Balakirev et al., 2015; Stingele et al., 2014).

**Figure 1:**
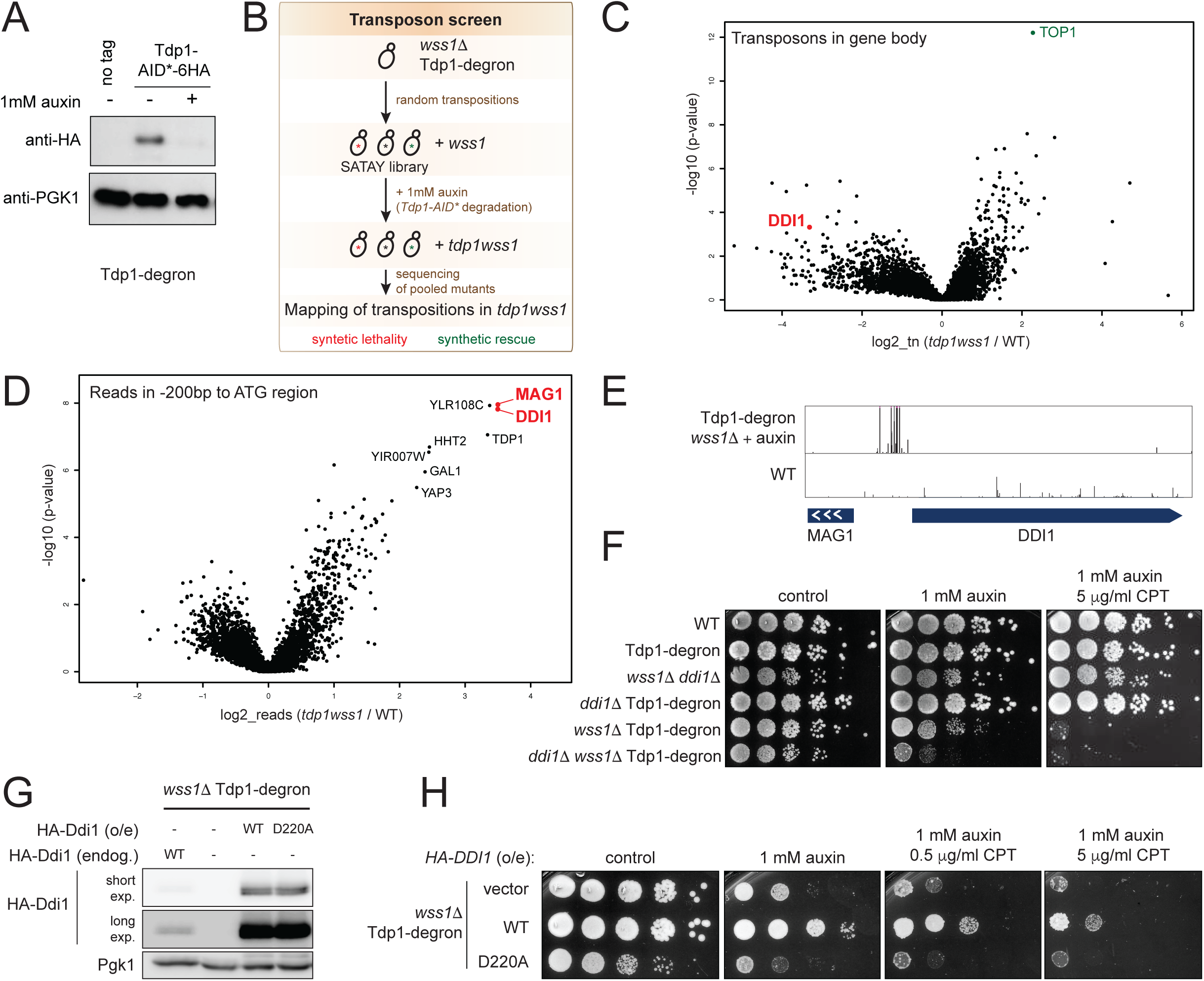
Yeast genetic screen of *tdp1wss1* identifies Ddi1 as another protease required for Top1cc processing. (A) Efficiency of Tdp1 depletion by the auxin-inducible degradation system. Tdp1-degron (*Tdp1-AID*-6HA pADH-TIR1*) and control strains were treated (+) or not (-) with 1 mM auxin for 8 h. Total cell extracts were subjected to immunoblotting. Pgk1 here and further was used as a loading control. (B) Outline of the SATAY transposon screen. (C-D) The transposon screen performed as illustrated in (B) identified the *DDI1* gene as one of the strongest negative genetic interactors of the *tdp1wss1* mutant. Transpositions in the mutant genome were compared to the pool of six other libraries. (C) Fold-change in the number of transposon per yeast gene bodies in the *tdp1wss1* library compared to a pool of unrelated libraries (log_2_, x-axis) and respective p-values (-log_10_, y-axis) are plotted. (D) Relative number of transposition events in the 200 bp region upstream of the ATG start codon (a presumable position of promoter). (E) Snapshot from the UCSC genome browser depicting transposon coverage of *DDI1* and *MAG1* genes in the mutant and one of the WT libraries. The height of each bar represents the number of reads for each detected transposon. Note that the height maxes out at 600. (F-H) Validation of the *DDI1* candidate gene from the screen. (F) *ddi1*Δ further negatively affects growth of *tdp1wss1*. (G-H) Elevated Ddi1 levels alleviate *tdp1wss1* phenotypes in protease-dependent manner. Serial 10x dilutions of cells were spotted on YEPD (F) or SC-leu (H) supplemented with 1 mM auxin and indicated amounts of camptothecin (CPT) and imaged after 48 h of growth. (G) HA-Ddi1 protein levels are compared by immunoblotting. *DDI1* was either HA-tagged at genomic locus (endog.), or *pADH-HA-DDI1* (WT) and *pADH-HA-ddi1-D220A* (catalytic protease mutant) were overexpressed from plasmids (o/e). See also Figure S1C for HA-Ddi1 protein levels in auxin and CPT.

The genes essential for Top1cc processing are expected to be synthetic lethal in the *tdp1wss1* screen. Among them, one of the most relevant non-essential ORFs crucial for *tdp1wss1* viability was *DDI1* (Figure 1C). The *DDI1* gene body was strongly depleted of transpositions, whereas the upstream region containing the promoter element was highly transposed (Figures 1D and 1E). Notably, this bidirectional promoter, driving the *DDI1* and *MAG1* divergent transcription, was the most frequently targeted in our screen, as compared to control libraries (Figure 1D). This can be reconciled assuming that insertion of a MiniDs transposon might be sufficient to overexpress a nearby gene. To test this hypothesis, we combined *tdp1wss1* mutant with *DDI1* deletion (Figure 1F) or overexpression from the strong *ADH* promoter (Figures 1G, 1H, and S1C). Indeed, whereas *ddi1*Δ further enhanced the *tdp1wss1* double mutant phenotype (Figure 1F), elevated levels of Ddi1 were sufficient to rescue the *tdp1wss1* growth defect and CPT sensitivity (Figure 1H), implying more efficient processing of Top1-DNA adducts. Strikingly, the putative Ddi1 protease activity was crucial for this rescue as the *ddi1-D220A* point mutant in the D[ST]G motif, a hallmark signature of aspartyl proteases, completely abolished the ability to suppress *tdp1wss1* (Figure 1H). Furthermore, *ddi1-D220A*, expressed at WT-like levels (Figure 1G), led to a dominant negative phenotype (Figure 1H), emphasizing the functional relevance of the conserved protease domain. Together, our results imply Ddi1-mediated proteolysis in the processing of persistent Top1ccs adducts.

### Processing of the Flp-nick induced crosslinks requires Ddi1

To gain insight into the molecular events accompanying DPC formation and metabolism, we used the previously described Flp-nick system mimicking Top1cc at a single genomic locus (Nielsen et al., 2009). Briefly, in this system the galactose-inducible *flp-H305L* recombinase mutant is targeted to the Flp recognition target (*FRT*) locus introduced artificially next to *ARS607*. An aberrant enzymatic reaction crosslinks the enzyme to DNA, leading to the formation of a DPC (Figure 2A). Similar to Top1cc, crosslinked Flp (Flp-cc) remains covalently bound to the DNA 3’ end through a tyrosine-phosphate bond and generates a single strand break on DNA. It was proposed previously that Top1 and Flp crosslinks have similar genetic requirements for repair (Nielsen et al., 2009), thus we first aimed to verify the importance of *TDP1, WSS1* and *DDI1* for cell survival in the presence of a single Flp-cc. As simultaneous deletion of these genes is nearly unviable due to Top1ccs toxicity, Flp-nick repair was examined in a *top1*Δ genetic background. Notably, one DPC lesion generated by Flp-nick at the *FRT* locus is insufficient to impair the growth of wild-type cells (Nielsen et al., 2009) and (Figure S2A). However, in the absence of Tdp1 and Wss1, the Flp-nick strain was sensitive to *flp-H305L* induction by galactose. Moreover, cell growth was further negatively affected by Ddi1 loss (Figures 2B, top panel, and S2A). Importantly, the most affected *ddi1*Δ*wss1*Δ*tdp1*Δ*top1*Δ mutant was sensitive to galactose only in the presence of the *FRT* site, supporting that Flp-cc formation is the underlying cause of this growth phenotype (Figure 2B, bottom panel).

**Figure 2:**
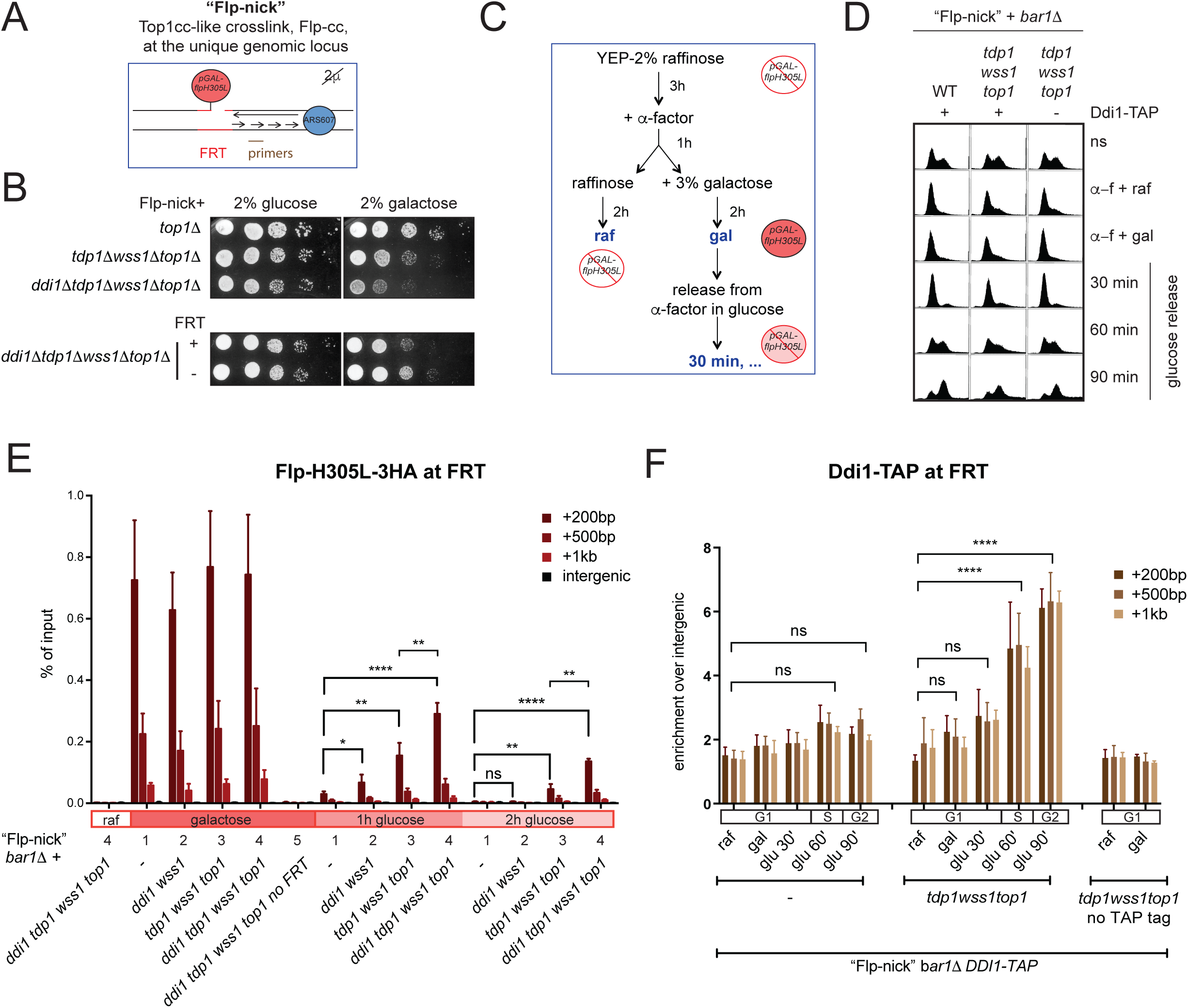
Ddi1 facilitates the repair of a Top1-like DNA-protein crosslink generated by mutant Flp recombinase at a single genomic locus. (A) Schematic representation of the Flp-nick system: the mutant galactose-inducible Flp-H305L recombinase forms a Top1cc-like DNA-protein crosslink when binds to the artificially introduced *FRT* site at the intergenic region next to *ARS607*. The system is based on the cir^0^ strain devoid of the naturally occurring 2μ plasmid that carries the *FLP* gene. (B) Top: the Flp-nick system is sensitive to the absence of Top1cc repair factors. Bottom: the growth defect of *ddi1*Δ*tdp1*Δ*wss1*Δ*top1*Δ depends on the presence of *FRT* binding site in the Flp-nick system. Flp-nick strains (cir^0^ *pGAL-flp-H305L tetO-FRT-ARS607*) combined with indicated deletions were pre-grown in YEP-2% raffinose and spotted on SC-glucose (control) or SC-galactose (Flp-nick induction) media. Images were taken after 60 h of growth. (C) Experimental design applied in Figures (D-F). (D) Cell cycle progression of Flp-nick strains was monitored by FACS analysis. ns, non-synchronized; α-f, alpha-factor; raf, raffinose; gal, galactose. (E) Loss of Top1cc repair factors, including Ddi1, delays the kinetics of Flp-cc removal from *FRT* locus. The levels of Flp-H305L-3HA covalently crosslinked to the *FRT* locus in the indicated deletion mutants were monitored by ChIP-qPCR without prior formaldehyde crosslinking. qPCR primers map downstream of FRT (+200 bp, +500 bp, +1 kb) as indicated in (A) and at non-related intergenic region. No Flp binding site (no *FRT*) and raffinose (no *pGAL-flpH305L-3HA* expression) samples were used as negative controls. Data are presented as the mean±SD for the percent of input of 3-4 independent replicates. To quantify the p-values, percent of input values for +200 bp were compared by 2-tailed T-test. (F) Ddi1-TAP is recruited to the Flp-bound *FRT* locus in an S-phase dependent manner after generation of Flp-cc. Ddi1-TAP occupancy close to the *FRT* locus monitored by ChIP-qPCR. The graph shows means±SD for n=3-5 of normalized to intergenic region qPCR signals; p-values for +200 bp were defined by 2-way ANOVA using Tukey’s multiple comparisons test.

We further asked whether the crosslinked Flp-H305L was retained longer on DNA in the absence of Tdp1, Wss1 and/or Ddi1. To address this question, an ectopically tagged *pGAL-flp-H305L-3HA* construct was induced with galactose in α-factor G1-synchronized cells, followed by simultaneous repression with glucose and further release from α-factor into the cell cycle (Figures 2C, 2D, and S2B). Flp retention at the *FRT* locus was monitored by chromatin immunoprecipitation (ChIP) without formaldehyde treatment. Upon galactose induction, Flp-H305L-3HA was recruited to *FRT* with a similar efficiency in different mutants, whereas the ChIP signal stayed at background levels in raffinose (no expression), as well as in the no *FRT* negative control (Figures 2E and S2C). Flp-induced DPC was rapidly evicted from the *FRT* locus in the strain with no additional mutations (Figure 2E, compare #1 in galactose, after 1h and 2h glucose release). In contrast, consistent with the defective growth on galactose, Flp-H305L-3HA was retained longer at the locus in the absence of Tdp1 and Wss1 (Figure 2E, #3). Notably, if Ddi1 was additionally depleted, the crosslinked Flp persisted even longer on the *FRT* locus (Figure 2E, compare *tdp1wss1top1 +/-DDI1*, #3 and #4 respectively), in line with the idea that Ddi1 participates in Flp-cc repair. We reasoned that Ddi1 would be recruited to the *FRT* locus if this protease directly participates in crosslink processing. Indeed, when monitoring Ddi1 recruitment to the *FRT*, a significant enrichment of the protein was found in proximity to the locus (Figure 2F). The Ddi1 signal was only detected if Flp-cc processing could not be successfully completed by Tdp1 and Wss1 (Figure 2F). Interestingly, Ddi1 recruitment to the *FRT* locus was observed only after 60 min of glucose repression, coinciding with the start of S phase (Figures 2D and 2F). These results indicate that when the conventional Top1cc repair factors fail to process Flp-cc, Ddi1 is recruited in a replication-dependent manner to facilitate DPC disassembly.

### Ddi1 functions redundantly with Wss1 in DNA-protein crosslink repair

The Tdp1 phosphodiesterase acts specifically on the DNA 3’-phosphate group covalently linked to a tyrosine residue, which limits Tdp1 to a specific subclass of DPCs (Connelly and Leach, 2004). In contrast, Wss1 has the capacity to proteolyze any DNA-bound protein (Stingele et al., 2014). We surmised that Ddi1 could also be active against a broad range of protein targets if functioning as a DPC protease. Interestingly, we noticed that endogenous Ddi1 levels were significantly elevated in *wss1*Δ, but not in the *tdp1* mutant (Figure 3A), suggesting that Ddi1 accumulation could represent a compensatory mechanism of Wss1 dysfunction. This observation prompted us to ask whether Ddi1 could have a general role in DPC processing.

**Figure 3:**
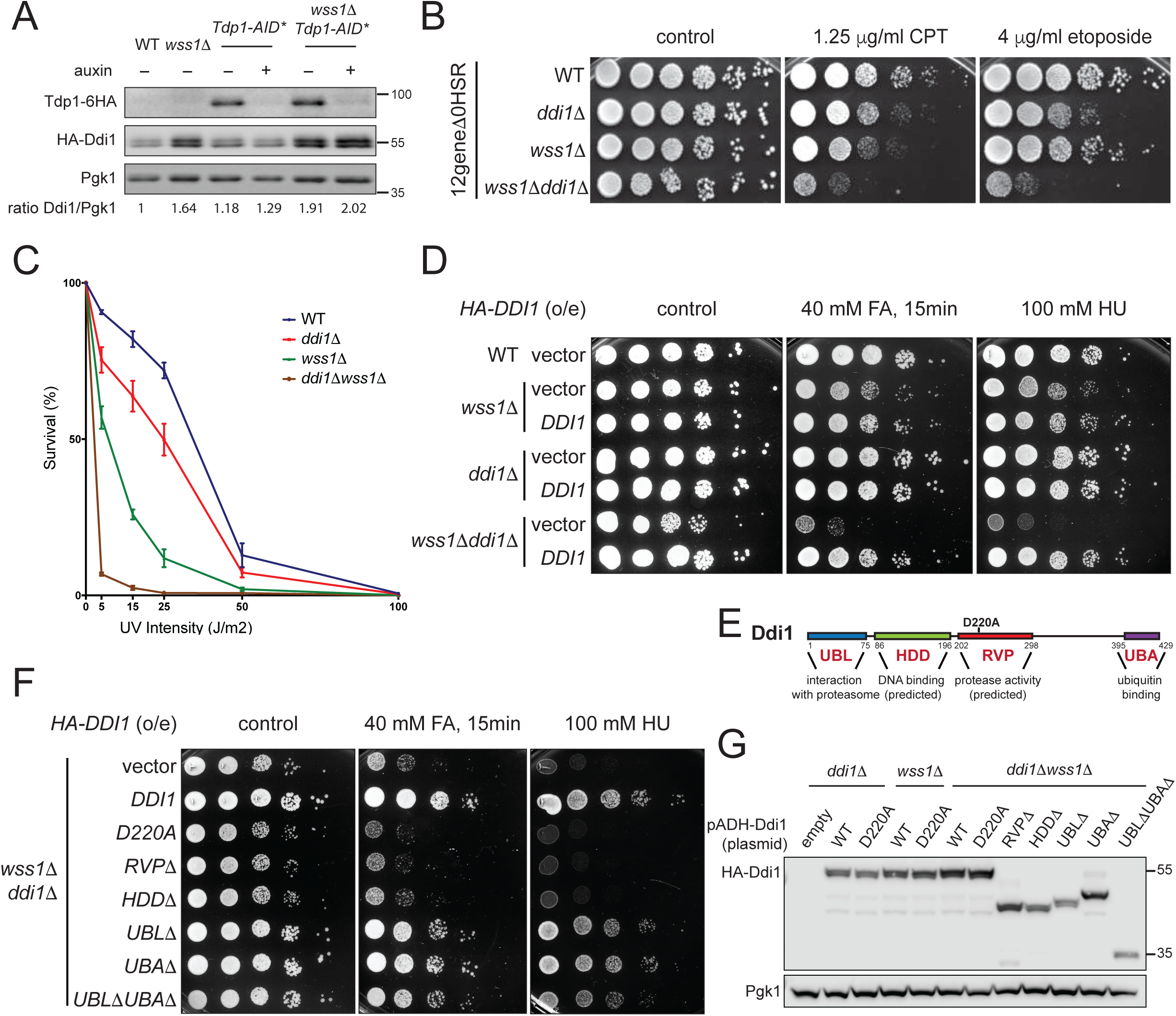
Ddi1 and Wss1 proteases provide resistance towards DPC-inducing agents and hydroxyurea. (A) Endogenous levels of Ddi1 are elevated in the absence of Wss1. Ddi1 was HA-tagged at the endogenous locus; total cell extracts were subjected to immunoblotting. Auxin was added for 6 h to degrade Tdp1-AID*-6HA. Quantifications of Ddi1/Pgk1 signals are shown at the bottom. (B) Ddi1 and Wss1 are redundantly required for the resistance towards enzymatic DPCs. The *12gene*Δ*0HSR* multitransporter mutant (Chinen et al., 2011) was used to reveal respective sensitivities to CPT (traps Top1cc) and etoposide (traps Top2cc) by spot test analysis. Images were taken after 72 h of growth. (C) *wss1*Δ and *ddi1*Δ mutants are sensitive to UV treatment. Cells were subjected to indicated doses of UV. Percent of survivals were normalized to non-treated samples for each genotype and plotted as means ±SEM for n=3. (D) Ddi1 is essential for survival of *wss1*Δ on formaldehyde (FA) and on hydroxyurea (HU). HA-Ddi1 was overexpressed from *pADH* promoter; cells grown in SC-leu were either pre-treated with 40 mM formaldehyde (middle) or spotted directly on 100 mM HU (right) and imaged after 48 h of growth. (E) Schematic representation of Ddi1 domain structure. (F) The catalytically active protease domain RVP and predicted DNA-binding domain HDD are essential for FA and HU tolerance. The experiment was performed as in (D). (G) Protein levels of overexpressed HA-tagged Ddi1 mutants used in (F) are compared by immunoblotting.

We first aimed to confirm the susceptibility of *ddi1*Δ mutant to enzymatically induced DPCs. The cell wall of conventional wild-type budding yeast strains is marginally permeable to drugs that enhance covalent trapping of topoisomerases (Nitiss and Wang, 1988). Consistently, despite the key role of Wss1 in DPC proteolysis, the loss of Wss1 does not sensitize yeast cells to Top1cc trapping drugs (Balakirev et al., 2015; Stingele et al., 2014). Likewise, in the W303 genetic background used in this study, *ddi1*Δ*tdp1* or *ddi1*Δ*wss1*Δ mutants are not sensitive to CPT, which induces Top1 adducts (Figure 1F). We thus took advantage of the *12gene*Δ*0HSR* multi-transporter mutant permeable for a broad range of chemical compounds (Chinen et al., 2011) to allow an efficient uptake of the topoisomerase trapping drugs. Both *ddi1*Δ or *wss1*Δ mutants showed a modest decrease in cell survival on CPT, which traps Top1ccs; remarkably, a strong synergistic effect was observed upon their simultaneous deletion (Figure 3B). The *12gene*Δ*0HSR* mutant was also used earlier to reveal the cytotoxicity of etoposide, a drug that traps topoisomerase 2 (Top2) in enzymatic DPCs (Wei et al., 2017). Notably, deletion of *DDI1* alone was sufficient to cause a growth defect in the presence of etoposide (Figure 3B). Similar to CPT, cell survival in presence of etoposide was greatly reduced in the *ddi1*Δ*wss1*Δ (Figure 3B).

To explore how Ddi1-depleted cells react to chemically derived DNA-protein crosslinks, we next induced DPC formation by UV light (Figure 3C) or reactive aldehydes (Figures 3D and 3F). We confirmed the previously observed *wss1*Δ hypersensitivity to UV light (O’Neill et al., 2004) and detected a mild effect of *ddi1*Δ on survival rates after UV exposure (Figure 3C). The survival of the UV-treated *ddi1*Δ*wss1*Δ mutant was additionally impaired (Figure 3C), similar to the synergistic phenotype observed in the presence of enzymatic DPCs. Formaldehyde (FA) is another potent crosslinking agent that non-specifically captures DPCs, and is therefore readily used to study the pathways involved in DPC repair (de Graaf et al., 2009). In contrast to severe FA sensitivity of mammalian cells lacking SPRTN protease, the Wss1 single mutant only weakly responds to FA treatment (Figure 3D) (Stingele et al., 2016; Stingele et al., 2014; Vaz et al., 2016). Strikingly, the combination of *ddi1*Δ and *wss1*Δ was hypersensitive towards FA, supporting the idea that Ddi1 could target a broad range of DPCs (Figure 3D).

Wss1 was previously described as a protein providing resistance towards hydroxyurea (HU) (O’Neill et al., 2004). Although HU does not crosslink proteins directly, it increases the *in vivo* production of reactive nitrogen (Jiang et al., 1997) and reactive oxygen species (Huang et al., 2016), both described to be an intrinsic cause of DPCs accumulation (Tretyakova et al., 2015). Analogous to DPC-trapping agents, *ddi1*Δ or *wss1*Δ mutants showed no or a weak sensitivity to HU respectively, while the absence of both proteases strongly sensitized cells to this drug (Figure 3D, right panel). Interestingly, CPT, but not HU sensitivity of *wss1*Δ and *ddi1*Δ*wss1*Δ strains could be rescued by *top1*Δ, supporting that Top1-induced crosslinks represent only a small subclass of DPCs targeted by Wss1 and Ddi1 (Figure S3A). Together, Ddi1 provides resistance towards different classes of DNA-protein induced crosslinks, especially in the absence of Wss1.

### Ddi1 function depends on its putative aspartic protease domain

Yeast Ddi1 is composed of 4 domains: ubiquitin-like (UBL), helical domain of Ddi1 (HDD), retroviral protease (RVP) and ubiquitin associated (UBA) domains (Figure 3E) (Gabriely et al., 2008). The UBL and UBA domains allow to classify Ddi1 as a proteasome shuttle, which physically connects the ubiquitinated cargo with the proteasome (Kaplun et al., 2005). These two domains were previously implicated in DNA damage-related functions of Ddi1, such as degradation of HO nuclease (Kaplun et al., 2005) or suppression of *pds1* checkpoint defects (Clarke et al., 2001). The RVP protease domain is the most conserved Ddi1 segment among eukaryotic orthologs, however its function remains the least understood. Structural analyses of the yeast and mammalian RVP domains strongly imply the presence of proteolytic activity, yet it has never been confirmed *in vitro* (Sirkis et al., 2006; Siva et al., 2016; Trempe et al., 2016). HDD is another evolutionally conserved domain (Siva et al., 2016) with potential for DNA recognition based on its weak similarities to DNA binding domains (Trempe et al., 2016). To test the importance of various domains for survival on genotoxins, we complemented *ddi1*Δ*wss1*Δ strain with Ddi1 domain mutants overexpressed from a strong *ADH* promoter. To our surprise, the ubiquitin receptor signature UBL/UBA domains were not required for survival on FA and HU (Figures 3F and 3G). In contrast, we observed the importance of HDD, probably designating a regulatory role for this domain. Strikingly, the RVP domain and the putative protease catalytic residue D220 were indispensable for *ddi1*Δ*wss1*Δ survival in the presence of drugs (Figures 3F and 3G). These data imply an essential role for Ddi1-mediated proteolysis, in agreement with the determinant role in Top1ccs tolerance (Figure 1H).

Previous screens of yeast and mammalian peptide libraries designed to detect putative Ddi1 protease activity *in vitro* revealed no Ddi1-dependent hydrolytic events (Siva et al., 2016; Trempe et al., 2016). Consistently, when we purified full-length or truncated versions of scDdi1 from bacterial or insect cells, we did not observe any cleavage of potential DNA-binding substrates such as Top1 or Histone 3 (Figure S3B and data not shown). *In vitro* cleavage reactions in the presence of potential activators such as 32nt or virion ssDNA, 32nt dsDNA, CPT, mono-or poly-Ub chains did not promote any obvious scDdi1-mediated proteolysis (Figure S3B and data not shown). Since our *in vivo* data strongly imply the Ddi1 protease activity in DPC resistance mechanisms, we concluded that *in vitro* cleavage requires the presence of other Ddi1 co-activators or co-factors.

### A functional proteasome accumulates on chromatin in the absence of Ddi1 and Wss1 proteases

The observations above support the idea that Ddi1 potential targets could be chromatin-associated proteins tightly bound or covalently crosslinked to DNA. Accordingly, in the absence of Wss1, the protease-dead Ddi1-D220A protein accumulates on chromatin, especially under stress conditions (HU or FA treatment) (Figure S4A). Interestingly, we noted that Ddi1 disfunction in *wss1*Δ background also triggers significant accumulation of the 26S proteasome complex on chromatin. Recently it was demonstrated that in *Xenopus* egg extracts 26S proteasome facilitates DPC proteolytic degradation redundantly with SPRTN (Larsen et al., 2018). To our knowledge, the yeast 26S proteasome has not been directly implicated in DPC repair, which prompted us to address this possibility. When the chromatin proteomes of *ddi1*Δ*wss1*Δ complemented with either *DDI1-WT* or *ddi1-D220A* grown in HU were compared by SILAC mass spectrometry (Figures 4A and S4B), we detected all 33 subunits of the 26S proteasome complex with a 1.3-2.65 fold increase in *ddi1-D220A* versus *DDI1-WT* (Figures 4B, 4C, and S4C).

**Figure 4:**
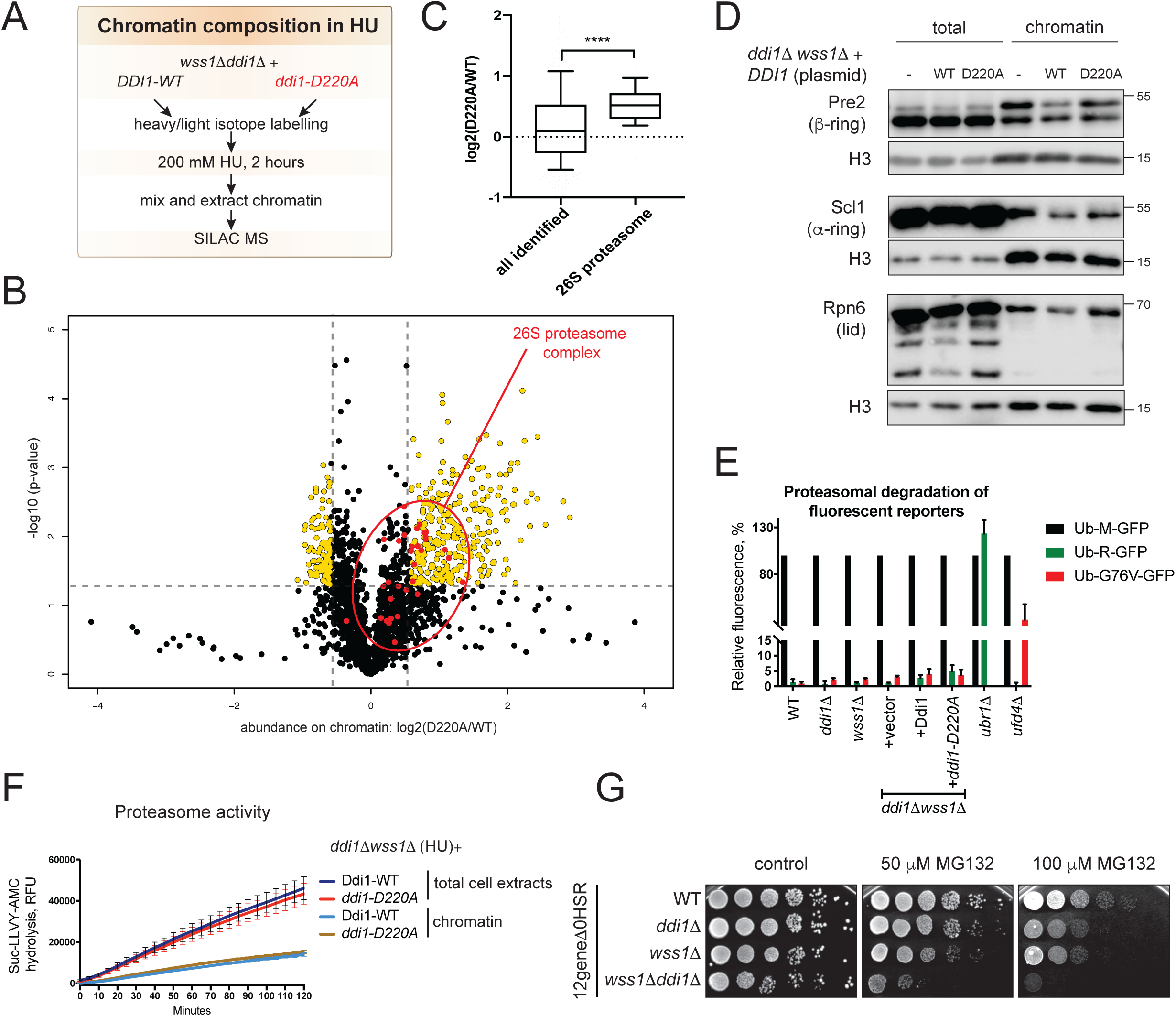
Functional 26S proteasome accumulates on chromatin in *ddi1*Δ*wss1*Δ in HU. (A) Experimental design used for the SILAC comparison of chromatin fractions between *ddi1*Δ*wss1*Δ complemented by overexpressed *HA-DDI1-WT* or *HA-*d*di1-D220A* grown in HU. Chromatin was isolated as in (Kubota et al., 2012). (B) Relative abundance of chromatin samples prepared as in (A). Three label-swap replicates were performed to quantify peptides by SILAC mass spectrometry. The volcano plot represents mean log_2_ (*ddi1*Δ*wss1*Δ*+HA-DDI1-WT* / *ddi1*Δ*wss1*Δ+HA-*ddi1-D220A*) intensity ratios for each quantified protein (x-axis) and their corresponding - log_10_ p-values (y-axis). Yellow dots represent proteins enriched at least 1.5 times with a p<0.05. Red dots depict components of the 26S proteasome complex. (C) The 26S proteasome is enriched on chromatin in *ddi1*Δ*wss1*Δ+*ddi1-D220A* mutant in HU. Data obtained in (B) were used to build a box and whiskers plot. The mean and 10-90 percentile are indicated. All identified proteins (2199) and 26S proteasome complex (33/33 components) were compared by Mann-Whitney test. (D) Validation of 26S proteasome subunits enrichment in *ddi1*Δ*wss1*Δ. HU treatment was performed as in (A). Three different TAP-tagged proteasome subunits were detected in total or chromatin fractions by immunoblotting with an anti-Protein A antibody. Histone H3 was used as a loading control. (E) No proteasomal disfunction is observed in single or combined *ddi1*Δ and *wss1*Δ mutants. Functional analysis of proteasome activity in live cells was monitored by flow cytometric analysis using various GFP reporters (Heessen et al., 2003). The control *ubr1*Δ and *ufd4*Δ mutants were used to reveal the degradation defects of Ub-R-GFP (N-end rule fluorescent substrate) or Ub-G76V-GFP (UFD signal substrate) respectively. The Ub-M-GFP reporter was used as a proteasome insensitive control. Where indicated, *ddi1*Δ is complemented with overexpressed *HA-DDI1-WT* or HA-*ddi1-D220A*. Data are represented as mean±SD of n=3. (F) The activity of total and chromatin-associated 26S proteasome is comparable in HU-treated *ddi1*Δ*wss1*Δ+*DDI1*-WT and *ddi1*Δ*wss1*Δ+*ddi1-D220A* samples prepared as in (A). Chymotrypsin-dependent hydrolysis of fluorogenic 20S proteasome substrate Suc-LLVY-AMC was used to measure proteasome activity in total extracts or in chromatin fractions; means±SEM of n=3 are plotted. 2-way ANOVA (Tukey’s multiple comparisons test) did not reveal significant changes between the mutants, with exception of control *ubr1*Δ and *ufd4*Δ. *ddi1*Δ and *wss1*Δ cells are sensitive to 26S proteasome inhibition. *12gene*Δ*0HSR* mutant was used to guarantee efficient uptake of MG132. Imaged were taken 72 h post-plating.

Next, we validated the accumulation of several proteasomal subunits on chromatin (Figure 4D). The proteasome could either be recruited to chromatin *via* a direct interaction with the UBL domain of Ddi1 (Saeki et al., 2002), or accumulate on chromatin independently of Ddi1 to compensate for the protease defective Ddi1-D220A. To distinguish between these two possibilities, we tested whether 26S proteasome subunits become enriched on chromatin in *ddi1*Δ*wss1*Δ complemented with an empty vector (excluding a direct interaction between Ddi1 and 26S proteasome). Although Ddi1 was absent in this mutant, we observed an enrichment similar to *ddi1-D220A* (Figure 4D), suggesting that proteasome recruitment to chromatin was not directly mediated by Ddi1. Thus, upon genotoxin-induced stress, the 26S proteasome accumulates on chromatin to compensate for loss of the Ddi1 protease independently of their direct interaction.

In *C. elegans* and mammalian cells, Ddi1 homologues regulate transcriptional induction of proteasomal subunit genes in response to stress (Koizumi et al., 2016; Lehrbach and Ruvkun, 2016). As these studies observe proteasome dysfunction in Ddi1 knockouts, we next asked whether the proteasome remains active in *ddi1*Δ*wss1*Δ cells. For that we measured the degradation efficiency of canonical N-end rule (Ub-R-GFP) and UFD (Ub-G76V-GFP) proteasome substrates (Heessen et al., 2003). In contrast to the control *ubr1*Δ and *ufd4*Δ mutants, no significant degradation defects of the GFP-based reporters were detected in *ddi1*Δ and/or *wss1*Δ mutants (Figure 4E). In addition, both chromatin and total cell extract samples derived from *DDI1-WT* or *ddi1-D220A* complemented cells showed similar 26S proteasome activity as defined by Suc-LLVY-AMC hydrolysis (Figure 4F). Last, both *ddi1*Δ or *wss1*Δ mutants were sensitive to proteasome inhibition by MG132 and *ddi1*Δ*wss1*Δ co-depletion had an additive phenotype (Figure 4G). Together, these data suggest that the 26S proteasome could compensate for Wss1 and Ddi1 and function as an independent protease, in agreement with the recent report about its role in DPC repair (Larsen et al., 2018).

### Ddi1 protease activity is required for the efficient turnover of the stalled Rpb1

RNA polymerases are central machineries involved in RNA production; DNA binding of these complexes is essential to guarantee efficient transcription. It is well described that the elongating RNA polymerase II permanently stalls in presence of a variety of DNA lesions (Beaudenon et al., 1999; Wilson et al., 2013; Woudstra et al., 2002). Arrested Rbp1, if not timely restarted, is subjected to ubiquitin-dependent degradation that depends on numerous factors, including the Cdc48 ATPase and the 26S proteasome (Wilson et al., 2013). Interestingly, when we selected from the SILAC chromatin proteomes described in Figure 4B a subgroup with the gene ontology term “DNA binding”, we detected 1.4x enrichment of the core RNA Pol II component Rpb1, encoded by the *RPO21* gene, in the *ddi1*Δ*wss1*Δ *+ ddi1-D220A vs ddi1*Δ*wss1*Δ *+ DDI1-WT* strain (Figures 5A and S5A).

**Figure 5:**
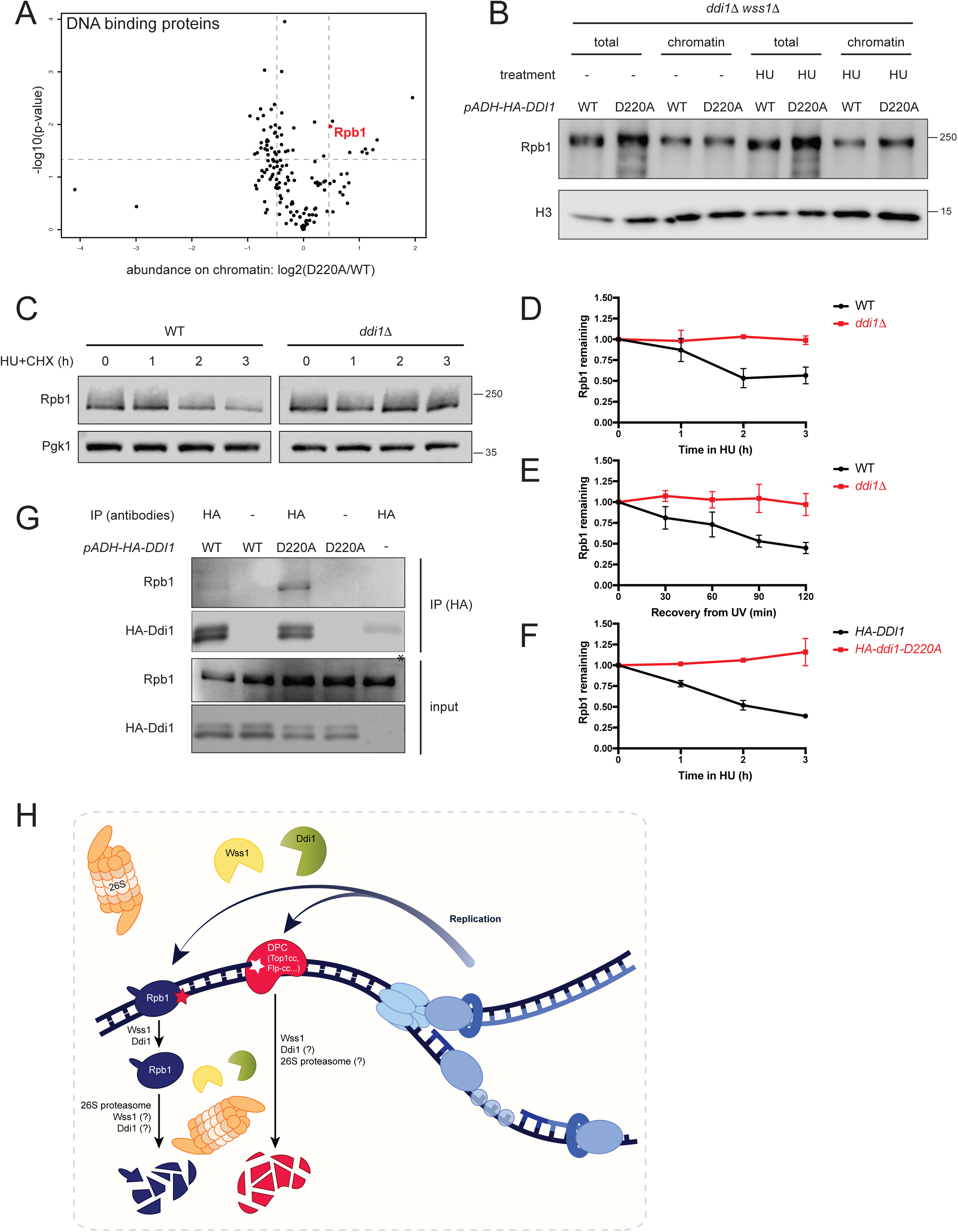
Ddi1 proteolytic activity is required for genotoxin-induced degradation of the core component of RNAPII complex Rpb1. (A) Rpb1 is one of the most significantly enriched DNA-binding proteins in the *ddi1*Δ*wss1*Δ+*ddi1-D220A* vs *ddi1*Δ*wss1*Δ+*DDI1* comparison. The volcano plot was generated using the subgroup of DNA-binding proteins found in the chromatin SILAC presented in Figure 4B. (B) Rpb1 accumulates on chromatin in *ddi1-D220A wss1*Δ treated for with 200 mM HU in the presence of cycloheximide (CHX, 100 μg/ml). Rpb1 levels in total and chromatin fractions were tested by immunoblotting. (C-F) Ddi1 by itself (C-E) and its protease activity (F) are required for the genotoxin-induced Rpb1 degradation. Rpb1 stability was examined in cells growing in the presence of 200 mM HU (C-D, F) or after 400 J/m^2^ of UV irradiation (E) in the presence of 100 μg/ml CHX at indicated time points. Rpb1 and Pgk1 levels in total cell extracts were probed by immunoblotting and quantified using fluorescent secondary antibodies. Relative Rpb1 to Pgk1 levels were set to 1 in the respective non-treated samples. Quantifications of at least 3 biological replicates are presented as means±SEM. (G) Ddi1 interact with Rpb1. Cells were UV-irradiated (400 J/m^2^) and HA-tagged overexpresses Ddi1 variants (-WT or -D220A) were immunoprecipitated with anti-HA antibodies; the Ddi1-Rpb1 association was analyzed by immunoblot.*, a signal from IgG heavy chains. (H) A model representing the crosstalk of yeast Ddi1, Wss1 and 26S proteasome in the proteolytic degradation of DPCs and Rpb1. Ddi1 and Wss1, probably in the context of replication, and the 26S proteasome are recruited to facilitate removal of DPC-trapped protein from DNA. Proteolytic disassembly of DPCs requires Wss1 (Stingele et al., 2014); Ddi1 and 26S proteasome are alternative candidate proteases which could directly facilitate DPC cleavage. Rpb1, stalled on damaged DNA templates (red star) also requires Ddi1 and Wss1 for efficient eviction from DNA and degradation.

Consistent with the mass spectrometry data, Rpb1 accumulates on chromatin upon HU-induced DNA damage in *ddi1*Δ*wss1*Δ *+ ddi1-D220A* cells (Figure 5B). We further asked whether Rpb1 stability could depend on Wss1 or Ddi1 proteases. Indeed, rapid decrease in total Rpb1 levels in response to genotoxin treatment could be detected in cells with cycloheximide-inhibited protein synthesis (Verma et al., 2011). Surprisingly, loss of Wss1 protease was sufficient to completely inhibit HU-induced degradation of Rpb1 (Figure S5B). Strikingly, Ddi1 was also absolutely required for the efficient genotoxin-induced Rpb1 turnover (Figures 5C, 5D, and 5F). The rapid degradation in the presence of HU typical for WT cells could not be detected in the *ddi1*Δ mutant (Figures 5C and 5D). Exposure to UV light resulted in the same Rpb1 degradation defect in *ddi1*Δ mutant (Figure 5E). Remarkably, proficient Rpb1 degradation was strongly dependent on the putative proteolytic activity of Ddi1, as the Ddi1-D220A mutant of the catalytic aspartate failed to support efficient degradation of Rpb1 in response to HU (Figure 5F). Last, we observed an interaction between Ddi1 and Rpb1 in co-immunoprecipitation experiments after UV exposure (Figure 5G), which was more prominent in the catalytically inactive *ddi1-D220A* mutant, suggesting that the protease may directly target Rpb1. Collectively, our data reveal that Wss1 and Ddi1, the two proteases that participate in DPC repair, also assist in the clearance of stalled core RNAPII subunits from DNA.

## DISCUSSION

### Interplay between the DPC-targeted proteases

Given the heterologous nature of crosslinking agents and the variety of proteins engaged in DNA transactions, the abundance of DPCs and their potential threat to genome integrity cannot be neglected. Deleterious consequences of these lesions stem from the bulky nature of a protein prone to crosslink formation, thus establishing the covalently bound protein as a central part of DPC. Albeit the protein moiety of a crosslink is the most evident target for repair, the existence of a proteasome-independent proteolytic pathway directed at DPCs was not addressed until recently (Duxin et al., 2014). This discovery was followed by the identification of the first DPC-dedicated proteases Wss1/SPRTN in different domains of life, including yeast, worms, mammals and plants (Enderle et al., 2019; Lopez-Mosqueda et al., 2016; Stingele et al., 2016; Stingele et al., 2014; Vaz et al., 2016). Up to date, the metalloproteases of the Wss1/SPRTN family remain the only enzymes which function is committed to DPC proteolysis. Interestingly, another putative ACRC protease, which shares the SprT-like domain with Wss1/SPRTN, was suggested as a candidate DPC protease in mammals, but the compelling experimental evidence has yet to be obtained (Fielden et al., 2018). Consequently, the pathways involved in proteolysis-mediated disassembly of DNA-protein crosslinks are only emerging and remain to be discovered.

This study identifies the yeast protease Ddi1 as a novel player in DPC repair. In contrast to the well-documented Wss1/SPRTN family of DPC-specific metalloproteases, Ddi1 belongs to a distinct class of retroviral aspartic proteolytic enzymes. In *Saccharomyces cerevisiae,* Wss1 and Ddi1 seem to function redundantly in DPC proteolysis, as supported by the observed synergistic effect of their co-depletion (Figures 1F, 3B-3D). Notably, *wss1*Δ alone neither has a deleterious growth phenotype, nor a strong sensitivity towards DPC-inducing agents (Balakirev et al., 2015; O’Neill et al., 2004; Stingele et al., 2014), consistent with the idea that the other DPC protease(s) could substitute Wss1 functions. In agreement with this model, the overexpression of Ddi1 in the presence of Top1cc or other DPC lesions is sufficient to partially suppress Wss1-related phenotypes (Figures 1F and 3D). Furthermore, cells lacking Wss1 accumulate the Ddi1 protein through an unknown mechanism (Figure 3A). It is worth noting that the susceptibility of *ddi1*Δ or *wss1*Δ cells towards DPC agents varies depending on the nature of the crosslinking agent (Figure 3), being in some cases more vulnerable in *wss1*Δ (FA, UV, CPT) or in *ddi1*Δ for others (etoposide). This can be explained by the potential diversity of mechanisms involved in the initial recognition of the lesions, thus defining the preferential recruitment of one or the other DPC protease.

The canonical ubiquitin-proteasome degradation pathway could also promote DPC disassembly independently of SPRTN (Larsen et al., 2018). Our observations indicate that Ddi1 also functions independently of the 26S proteasome, despite the previously described proteasome shuttle role (Figure 4). Additionally, the proteolytic activity of the 26S proteasome complex could explain the visible removal of crosslinked Flp-H305L from the *FRT* locus in the absence of Wss1 and Ddi1 proteases (Figure 2E). In agreement with the redundant role of the three pathways, 26S proteasome components were previously found among top negative genetic interactors for both Ddi1 and Wss1 (Costanzo et al., 2016). Moreover, the *ddi1*Δ*wss1*Δ mutant accumulates 26S proteasome on chromatin (Figure S4D) and is sensitive to the MG132 treatment in the absence of any genotoxins (Figure 4G). Wss1 and Ddi1 dysfunction probably leads to accumulation of endogenous lesions, thus triggering 26S proteasome recruitment as a feedback response.

Wss1/SPRTN partners with the Cdc48/p97 segregase (Balakirev et al., 2015; Mosbech et al., 2012; Stingele et al., 2014) and in yeast this interaction was previously shown to be indispensable for Wss1 function in Top1cc repair (Stingele et al., 2014). Interestingly, recent co-immunoprecipitation experiments coupled with mass spectrometry revealed that Ddi1 also interacts with Cdc48, however the study didn’t investigate whether it also occurs on chromatin (Kama et al., 2018). We speculate that Ddi1-Cdc48 interaction could possibly play a role in the extraction of crosslinked protein from the lesion *via* a mechanism similar to Wss1 (Stingele et al., 2014).

### Ddi1: a novel DPC dedicated protease?

Our *in vivo* experiments with the putative catalytic aspartate residue mutation *ddi1-D220A* strongly implicate the importance of Ddi1 predicted proteolytic activity in DPC-related functions. For example, *ddi1-D220A* has deleterious phenotypes in the presence of crosslinking agents (Figures 1H and 3F) and exhibits defective Rpb1 turnover (Figure 5F). Another line of evidence that supports the essential role of Ddi1-mediated proteolysis comes from the observed dominant negative phenotypes of the overexpressed *ddi1-D220A* (Figure 1H). The inactive protease is expected to compete for substrates with functional Ddi1 and Wss1 and to increase the interaction time with its hypothetical targets. In accordance with this, the catalytically dead Ddi1-D220A mutant accumulates on chromatin upon stress (Figure S4A) and interacts with Rpb1 more efficiently than Ddi1-WT (Figure 5G), possibly as a result of the blocked enzymatic reaction. Taken together, we speculate that Ddi1 can directly facilitate DPC proteolysis.

Supporting our hypothesis, structural studies revealed that the Ddi1 protease domain has a big cavity allowing it to accommodate large substrates (Sirkis et al., 2006; Siva et al., 2016; Trempe et al., 2016). Such a configuration of the active site should be perfectly suited for the unspecific cleavage of bulky proteins trapped in DPCs. However, to date, the biochemical evidence of *in vitro* hydrolysis catalyzed by Ddi1 are lacking so far. Despite our attempts, we also could not detect the proteolytic activity of purified Ddi1, consistent with previous reports (Siva et al., 2016; Trempe et al., 2016). This may not be so surprising, as the DPC protease is expected to be tightly regulated in order to restrain uncontrolled proteolysis of unwanted substrates. We believe that Ddi1 is activated in response to some signal(s), however the nature of these stimuli remains to be discovered.

We identified the RNA polymerase II core subunit Rpb1 as one potential substrate of Wss1 and Ddi1 proteases. Stalled Rpb1 was previously described as a canonical substrate for both Cdc48 and 26S proteasome (Beaudenon et al., 1999; Verma et al., 2011; Wilson et al., 2013). Rpb1 degradation in response to genotoxin treatment is a defensive mechanism, which allows cells to restart transcription otherwise blocked by RNA Pol II collisions with genotoxin-induced lesions (Wilson et al., 2013). Defective degradation of Rpb1 in *ddi1*Δ and *wss1*Δ mutants and the importance of Ddi1 protease activity can be interpreted as a direct action of these enzymes on stalled Rpb1. In support of this hypothesis, SPRTN cleaves with a similar efficiency free and a DPC-trapped topoisomerases (Vaz et al., 2016), suggesting that tightly bound non-crosslinked proteins could also be substrates for a DPC protease. Thus, we speculate that these observations reflect a direct effect of Ddi1 and Wss1 on Rpb1 degradation.

### How could Ddi1 be recruited and activated?

One outstanding question concerns the potential activation mechanism and the mode of Ddi1 recruitment to DPC lesions. We detected Ddi1 at the site of DPC formation and the signal appears in an S-phase dependent manner (Figure 2F). We only observe a significant enrichment of Ddi1 at the locus if other pathways that facilitate crosslink repair are inactivated (Wss1 and Tdp1 in case of Flp-cc). Thus, a persistent crosslink in S-phase could be a signal for Ddi1 recruitment. It is worth mentioning that the yeast Flp recombinase expressed from yeast 2µ circles could be modified by SUMO to prevent DNA damage accumulation (Xiong et al., 2009). Wss1, on the other hand, is able to recognize and bind this posttranslational modification *via* two canonical SIM domains (Stingele et al., 2014) and Wss1-mediated proteolysis is activated by high molecular weight SUMO species (Balakirev et al., 2015), which can explain the preference of Wss1 for Flp-ccs. In this scenario Wss1 could be targeted to SUMO-modified crosslinks, whereas Ddi1 recruitment would be a secondary event triggered by another signal, e.g. ubiquitin. This order of events could be quite specific to Flp-ccs and distinct initial signals may target Ddi1 to other types of DPCs in a Wss1-independent manner.

The Ddi1 domain structure reveals a typical signature of ubiquitin-binding motifs. Furthermore, multiple studies demonstrated strong mono-Ub and poly-Ub binding affinity of both UBL and UBA domains *in vitro* (Bertolaet et al., 2001; Nowicka et al., 2015; Siva et al., 2016; Trempe et al., 2016). In our hands, ubiquitin chains do not activate proteolytic activity of purified full-length Ddi1 (Figure S3B), however we cannot exclude that the concentration or the type of Ub chains were not optimal in our reactions. Alternatively, the presence of free ubiquitin may not be sufficient, as its conjugation to the substrate could be essential to promote Ddi1 activity. In addition to activation, ubiquitin-binding domains could play a role in substrate recognition. Although we observed no evident role for UBL and UBA in cell growth on genotoxins (Figure 3F), in these experiments we used overexpression of Ddi1 mutants, which could mask the contribution of these domains. In contrast, *ddi1-UBL*Δ or *ddi1-UBA*Δ mutants expressed endogenously suggested the importance of both domains; however this may be explained by a compromised protein level of the corresponding Ddi1 mutants (Figures S3C and S3D). Thus, future studies are required to reveal the importance of ubiquitin in Ddi1 recruitment and activation.

In addition to ubiquitin binding, Ddi1 also has *in vitro* affinity for Rub1 (yeast NEDD8 homologue) and Rub1-Ub heterodimers (Singh et al., 2012), but the functional significance of these interactions is elusive. Furthermore, Ddi1 is phosphorylated on S341, T346 and T348 residues within the Sso1 binding domain (Gabriely et al., 2008). Many aspartic proteases, including HIV protease, are expressed as zymogens and are activated by (auto-) proteolysis (Eder et al., 2007). Likewise, Ddi1 is a substrate for metacaspases (Bouvier et al., 2018) and contains a PEST sequence that typically marks proteins for degradation (Gabriely et al., 2008). However, neither phosphorylation nor PEST sites are essential for Ddi1 HU and FA resistance (Figures S3C and S3D), but we cannot exclude their secondary role in Ddi1 regulation.

### Could Ddi1 have an evolutionary conserved role in DPC repair?

*DDI1* was duplicated during evolution yielding two human *DDI1* and *DDI2* genes. The duplication was preceded by the loss of the C-terminal UBA domain in the vertebrate lineage (Siva et al., 2016), emphasizing the evolutionary relevance of the central protease domain of Ddi1, rather than the UBA ubiquitin receptor signature of this protein. Interestingly, in higher eukaryotes, Ddi1 homologues activate the SKN-1/Nrf1 transcription factor in a protease-dependent manner (Koizumi et al., 2016; Lehrbach and Ruvkun, 2016). This probably represents an evolutionary innovation, as SKN-1/Nrf1, the master regulator of 26S proteasome transcription, is not conserved in yeast. On the other hand, hDDI1 and hDDI2 were recently detected on replication forks where they facilitate the protein barrier removal; however, the importance of hDdi1/2 protease activity was not addressed in this study (Kottemann et al., 2017). Given the tight link between 26S proteasome-mediated proteolysis and DPC repair, it is possible that the two aforementioned functions of human Ddi1 homologues co-evolved in higher eukaryotes. As the current study implicates yeast Ddi1 in the control of DPC proteolysis, it would be interesting to address in the future whether DNA-protein crosslinks are the biological targets of the DDI1 and DDI2 proteases in higher eukaryotes.

## Supporting information

Supplementary table 1. List of strains and plasmids

Supplementary data 1. Data analysis and results of SILAC mass spectrometry related to Figure 4.

## ACKNOWLEDGEMENTS

We thank Helle Ulrich for providing auxin degron plasmids, Jeffrey Gerst and Jean-Francois Trempe for *DDI1* constructs, Lotte Bjergbaek for the Flp-nick system, Benjamin Albert for the *top1*Δ and Takeo Usui for the *12gene*Δ*0HSR* strains. We thank Margot Riggi for the graphical work. We are grateful to Geraldine Silvano for technical assistance; Lionel Jond and Sandra Citi for the help with insect cells; Kaushik Bhattacharya for the help with proteasomal activity assays; to Maksym Shyian, Vincent Dion, Thanos Halazonetis and all members of Stutz laboratory for critical reading of the manuscript, their comments, discussions and useful suggestions. This work was supported by funds from the Swiss National Science Foundation (grants 31003A 153331 and 31003A_182344 to FS), iGE3, the Canton of Geneva, the NCCR Chemical Biology (R.L. and M.P) and ERC TORCH (R.L.).

## AUTHOR CONTRIBUTIONS

Conceptualization: N.S. Methodology: N.S., A.N., I.B., M.P., A.H.M., R.L., B.K., F.S. Investigation: N.S., A.N., I.B., A.H.M., M.P. Data Curation: N.S., B.K., A.H.M., M.P., A.N. Writing – Original Draft: N.S. Writing – Review and Editing: N.S., F.S., M.P., B.K., A.N., I.B. Supervision: F.S., B.K., R.L. Funding acquisition: F.S.

## STAR Methods

### CONTACT FOR REAGENT AND RESOURCE SHARING

Further information and requests for resources and reagents should be directed to and will be fulfilled by the Lead Contact, Françoise Stutz

### EXPERIMENTAL MODEL AND SUBJECT DETAILS

#### Saccharomyces cerevisiae

##### Yeast strains

*Saccharomyces cerevisiae* strains were derived from W303 [*leu2-3,112 trp1-1 can1-100 ura3-1 ade2-1 his3-11,15*] or S288C (BY4741 or BY4742) [*his3*Δ*1 leu2*Δ*0 lys2*Δ*0 met15*Δ*0 ura3*Δ*0*] genetic backgrounds. A complete list of yeast strains is available in Table S1.

##### Growth medium

All yeast cultures were grown at 30°C. Liquid YEP-or SC-based media were supplemented with 2% glucose or other sources of sugar (2% raffinose or 2-3% galactose). SC-or YEPD-based plates were supplemented with 20 g/l agar. To select against URA3 marker 1 mg/ml of 5*-*Fluoroorotic Acid was added. Selection for dominant markers was performed on YEPD-based medium supplemented with 200 μg/ml G418, 200 μg/ml clonNAT or 50 μg/ml Hygromycin B.

YEPD medium: 1% yeast extract, 2% peptone, 2% glucose.

SC medium: 1.7 g/l yeast nitrogen base, 5 g/l ammonium sulfate, 2% agar, 2% glucose, 0.87 g/l dropout mix. Dropout mix composition: 0.8 g adenine, 0.8 g uracil, 0.8 g tryptophan, 0.8 g histidine, 0.8 g arginine, 0.8 g methionine, 1.2 g tyrosine, 2.4 g leucine, 1.2 g lysine, 2 g phenylalanine, 8 g threonine.

SD+2% galactose-adenine agar plates used for the transposon screen: 1.7 g/l yeast nitrogen base, 5 g/l ammonium sulfate, 2% agar, 2% galactose, 0.03 g/l isoleucine, 0.15 g/l valine, 0.02 g/l arginine, 0.02 g/l histidine, 0.1 g/l leucine, 0.03 g/l lysine, 0.02 g/l methionine, 0.05 g/l phenylalanine, 0.2 g/l threonine, 0.04 g/l tryptophan, 0.03 g/l tyrosine, 0.02 gl/ uracil, 0.1 g/l glutamate, 0.1 g/l aspartate.

#### Insect cells

SF9 insect cells were grown at 27°C in liquid TC-100 medium supplemented with Penicillin/Streptomycin, FBS and CD lipid concentrate.

#### Bacteria

DH5α *E. coli* strains were grown in LB medium at 37°C. For plasmid selection LB-2% agar plates were supplemented with 50 μg/ml of ampicillin.

## METHOD DETAILS

### Yeast techniques

#### Yeast transformation

Log phase yeast cultures were resuspended in LiTE buffer (100 mM LiAc, 10 mM Tris pH 7.5, 1 mM EDTA) and mixed with 100 μg/ml salmon sperm ssDNA, 37.28% [w/v] PEG4000 and DNA (200 ng of pDNA or purified PCR fragment). Cells were incubated for 1 h at 30°C, then supplemented with 6% DMSO and a heat shock was performed for 10 min at 42°C. Single colonies were isolated from selective media.

#### Genetic crosses

To obtain a combination of mutants, yeast strains of opposite mating type were mixed and grown overnight on rich YEPD medium, transferred on different media to select for prototrophic diploids and sporulated for 4-5 days on KAC plates (20 g/l potassium acetate, 2.2 g/l yeast extract, 0.5 g/l glucose, 0.87 g/l dropout mix, 20 g/l agar, pH 7.0) to obtain tetrads. Asci were digested with 0.5 mg/ml zymolyase 20T to release spores that were separated by dissection to obtain the desired genotypes.

#### Yeast colony PCR

Yeast strains were genotyped by direct resuspension in Phire Green Hot Start II PCR Master Mix supplemented with oligonucleotides according to manufacturer recommendations.

#### Construction of strains

Single mutations in yeast genomes were introduced by transformation. Oligonucleotides and template plasmid DNA used to generate PCR products for homologous recombination are listed in Supplementary Table 1. Correct modifications of genomic DNA were verified by colony PCRs and/or sequencing. An epitope tag insertion was additionally checked by immunoblotting. Mutants with multiple genomic modifications were obtained by genetic crosses.

#### Spot test

Log phase yeast cultures were diluted to OD_600_=1-1.5, five 10x serial dilutions were prepared in a sterile 96 well plate and spotted on YEP-or SC-based agar plates supplemented with auxin, camptothecin (CPT), hydroxyurea (HU), etoposide or MG132. Formaldehyde (FA) treatment was performed in 1 ml yeast liquid cultures for 15 min, cells were washed twice with 1 ml H2O and spotted on plates without additional drugs.

#### UV irradiation and quantification of survival

Cultures were diluted to desired concentrations and plated on YEPD in triplicate for each UV dose and dilution. Plates were dried for 15 min at room temperature before irradiation with indicated doses of UV using Stratalinker 2400. After irradiation, plates were grown in the dark for 2 days and colonies were counted to define percent of survivals.

#### Flp-nick induction

Induction of the *pGAL-flp-H305L* construct in *FRT*-containing strains (Flp-nick) was essentially performed as described in (Nielsen et al., 2009). All Flp-nick strains used for ChIP experiments contained the *bar1*Δ mutation to facilitate efficient G1 arrest. Cells in log phase were grown in YEP-2% raffinose and synchronized for 1h with 200 ng/ml α-factor prior to *pGAL-flp-H305L* induction with 3% galactose. Half the amount of α-factor was re-added together with galactose. To repress *pGAL-flp-H305L*, cells were washed twice with YEP no sugar medium and resuspended in YEP-2% glucose. 1 ml aliquots were collected at different time-points to monitor cell cycle progression by FACS (see below).

### SATAY library generation

The SATAY transposon screen was essentially performed as described (Michel et al., 2017). Two independent clones of the *tdp1wss1* mutant in BY4741 genetic background [*wss1Δ::HIS3 leu2Δ0::TIR1-LEU2 Tdp1-AID*-6HA::HPH ade2Δ::KANMX6 MATa*] were transformed with pBK257. Transformants were pre-grown in SC-ura+2% raffinose/0.2% glucose to saturation and plated on SD+2%galactose-adenine agar plates and grown for 3 weeks to generate a library containing approximately 3.7 million clones (estimated from counting 1 cm^2^ area for 10 different plates). Collected cells were pooled and diluted to OD_600_=0.45 in 2 liters of SC+2%glucose-adenine medium and grown overnight until saturation. The library was re-diluted to OD_600_=0.1 and 1 mM auxin was added; after 24 h of growth cells were re-diluted to OD_600_=0.1 in a fresh auxin-containing medium and grown to saturation (∼46 h).

Genomic DNA was extracted, digested with DpnII and NlaIII, circularized and PCRs were performed with P5_MiniDs and MiniDs_P7 oligos and quantified by absorbance at 260 nm as detailed in (Michel et al., 2017). Equal amounts of purified PCR products obtained from DpnII-and NlaIII-digested DNA were pooled and sequenced using MiSeq v3 chemistry adding 3.4μl of 100 μM 688_minidsSEQ1210 primer according to manufacturer recommendations.

Bioinformatics analysis was performed essentially as described in Michel et al. 2017, and the coordinate and number of associated sequencing reads were saved as .bed and .wig file for upload into a genome browser.

### Protein extraction

Exponentially growing yeast cultures corresponding to 5-10 OD_600_ were fixed with 6.25% of trichloroacetic acid, kept on ice for 10 min, pelleted, washed twice with 100% acetone and dried under vacuum. Dry pellets were resuspended in urea buffer (50 mM Tris-Cl pH 7.5, 5 mM EDTA, 6 M Urea, 1% SDS), mixed with 200 μl of 0.5 mm glass beads and homogenized in MagNA Lyser 5 times for 45 s at 4°C. Solubilization was performed for 10 min at 65°C followed by 10 min centrifugation at 18000 g. Proteins were resuspended in 1.5x volume of sample buffer (3% SDS, 15% glycerol, 0.1 M Tris pH 6.8, 0.0133% bromophenol blue, 0.95 M 2-mercaptoethanol) and boiled for 10 min.

### Immunoblotting

Proteins were resolved by SDS-PAGE using 6-14% gels, transferred to nitrocellulose membrane, blocked with 5% dry milk dissolved in TBS-T (150 mM NaCl, 20 mM Tris-HCl, 0.05% Tween, pH 7.4), followed by incubation with primary and HRP-coupled secondary antibodies diluted in TBS-T. Signal was revealed with WesternBright HRP substrate; images were taken with Li-COR Odyssey Imaging System and quantified using Image studio v.3.1. IRDye fluorescent secondary antibodies were used to quantify relative Rpb1 levels.

### Co-immunoprecipitation

The protocol of co-IP experiments is based on (Verma et al., 2011). 100 ml of yeast cells (OD_600_=1-1.5) were UV-irradiated (400 J/cm2), washed twice with PBS (137 mM NaCl, 2.7 mM KCl, 10 mM Na_2_HPO_4_, KH_2_PO_4_, pH 7.4) and collected in the dark. Pellets were resuspended in 1 ml of Lysis buffer C (50 mM HEPES, pH 7.6, 1.0% Triton, 0.1% sodium deoxycholate, 0.5 M potassium acetate, 10% glycerol, 2 mM EDTA, 2 mM EGTA, 1 mM PMSF, 50 mM β-glycerophosphate, 2 mM NaF, 25 mM NEM, Protease Inhibitor cocktail), mixed with 500 μl of glass beads and homogenized by MagNA Lyser, 5x 45 s cycles of 6500 rpm, cooling samples at 4°C for at least 1 min in between the cycles. All subsequent procedures were performed on ice. Lysates were clarified by 25 min centrifugation at 18000 g, 30 μl input was collected and the remaining lysates were mixed with 1 μl anti-HA antibody. Immunoprecipitations were incubated overnight with rotation at 4°C; 25 μl Protein G Dynabeads in Lysis C buffer were then added to each lysate and samples were rotated for 1 h. Dynabeads containing protein complexes were washed 6 times with Wash buffer D (25 mM HEPES, pH 7.5, 100 mM potassium acetate, 150 mM KCl, 0.25% Tween 20, 10 % glycerol, 50 mM β-glycerophosphate) and boiled in 30 μl 1x sample buffer. Input and IP samples were further subjected to immunoblotting with anti-HA and anti-Rpb1 antibodies.

### SILAC mass spectrometry to compare chromatin proteomes

#### Isotopic labelling and mixing of cultures

Yeast cell cultures [*lys1Δ; arg4Δ*] were grown in 125 ml of SC-leu-arg-lys medium supplemented with “heavy” (50 mg/l of Arg-10 and Lys-8) or “light” (50 mg/l of Arg-0 and Lys-0) amino acids mixes for at least 8 doubling cycles before drug treatment. 200 mM HU was added for 2 h and cultures were collected at OD_600_=0.8-1.2. Equivalent numbers of differently labelled cells were mixed for each replicate; for replicates 1 and 3 *DDI1-WT* cultures were labelled with “light” and the *ddi1-D220A* mutant with “heavy” amino acids; for the replicate 2 was labelling conditions were swapped. 10 ml of each culture were collected to analyze the labelling efficiency, which was estimated to be 97%. Mixed cultures were washed with 20 ml of 1x PBS, the supernatant was removed completely and pellets were frozen in liquid nitrogen.

#### Isolation of chromatin fractions

Chromatin fractions were isolated as in (Kubota et al., 2012). Pellets were resuspended in 10 ml of pre-spheroplasting buffer (100 mM PIPES/KOH, pH 9.4, 10 mM DTT, 0.1% sodium azide) and incubated 10 min at room temperature. After 3 min centrifugation at 1800 g, pellets were resuspend in 10 ml of spheroplasting buffer (50 mM KH_2_PO_4_/K_2_HPO_4_, pH 7.4, 0.6 M Sorbitol, 10 mM DTT, 0.5 mg/ml Zymolyase-20T and 5% Glusulase) and incubated for 30 min at 37°C. Spheroplasts were pelleted for 3 min at 1800 g at 4°C and washed twice with 5 ml of ice-cold Wash buffer (20 mM KH2PO4/K2HPO4, pH 6.5, 0.6 M Sorbitol, 1 mM MgCl2, 1 mM DTT, 20 mM β-glycerophosphate, 1 mM PMSF, 1 protease inhibitor tablet/10 ml), resuspended in 5 ml fresh Wash buffer, overlaid on 5 ml of 7.5% Ficoll–Sorbitol cushion buffer (7.5% Ficoll, 20 mM KH_2_PO_4_/K_2_HPO_4_, pH 6.5, 0.6 M Sorbitol, 1 mM MgCl_2_, 1 mM DTT, 20 mM β-glycerophosphate, 1 mM PMSF, 1 protease inhibitor tablet/10 ml) and centrifuged at 4000 g for 5 min at 4°C. Tube walls were washed with 10 ml of ice-cold Wash buffer, then pellets were resuspended in 200 μl of ice-cold Wash buffer. To obtain the whole cell extract (WCE) fraction, 5 μl of the suspension was transferred into a new tube and resuspended in 50 μl of EBX buffer (50 mM HEPES/KOH, pH 7.5, 100 mM KCl, 2.5 mM MgCl_2_, 0.1 mM ZnSO_4_, 2 mM NaF, 0.5 mM spermidine, 1 mM DTT, 20 mM β-glycerophosphate, 0.25% Triton X-100, mM PMSF, 1 protease inhibitor tablet/10 ml). The remaining 195 μl were mixed with 14 ml of 18% Ficoll buffer (18% Ficoll, 20 mM KH_2_PO_4_/ K2HPO_4_, pH 6.5, 1 mM MgCl_2_, 1 mM DTT, 0.01% NP40, 20 mM β-glycerophosphate, 1 mM PMSF, 1 protease inhibitor tablet/10 ml) and resuspended by magnetic stirring and 10 strokes in Potter-Elvehjem homogenizer. Lysed cells were centrifuged twice at 5000 g for 5 min at 4°C; the supernatant was retained at this stage. To pellet nuclei, the solutions were centrifuged at 16100 g for 20 min and the supernatants containing the cytoplasmic fractions (C) were collected. Sides of the tubes were washed with 1 ml of cold Wash buffer that was aspirated without further spin; the pellets were resuspended in 1 ml of Wash buffer and centrifuged for 5 min at +4°C at 4000 g. Nuclei were resuspended in 200 μl of EBX buffer and incubated on ice for 10 min. The lysates were overlaid on EBX containing 30% sucrose and chromatin was pelleted at 16000 g for 10 min at 4°C. The sucrose layer was discarded, sides of the tube were washed with 1 ml of EBX buffer which was immediately aspirated. Chromatin pellets were resuspended in 1 ml of EBX buffer and centrifuged at 10000 g for 2 min. In order to remove all residues of detergents and inhibitors, chromatin pellets were additionally washed three times in 500 μl of 50 mM ammonium bicarbonate, every time pelleted by 10000 g centrifugation for 2 min. For the last wash, the suspension was transferred to the nonstick tubes, 400 μl of supernatant were removed after centrifugation, the sides of the microcentrifuge tubes were washed and samples were centrifuged again for 5 min. The supernatants were removed completely and pellets were frozen at −20°C until further processing. WCEs, aliquots of cytoplasmic (C) and chromatin pellet (Ch) fractions were mixed with sample buffer and boiled for 10 min for qualitative analysis by immunoblotting.

#### Peptide preparation and mass spectrometry

Chromatin pellets were resuspended in 8 M urea in 50 mM ammonium bicarbonate (ABC). Samples corresponding to 30 μg of protein quantified by Bio-Rad protein assay were incubated with 5 mM Tris (2-carboxyethyl) phosphine and 5 mM iodoacetamide for 30 min at room temperature in the dark. After addition of dithiothreitol to 5 mM, samples were incubated for an additional 30 min at room temperature. The concentration of urea was diluted to 5.5 M with 50 mM ABC and MgCl_2_ was added to 2 mM. The samples were incubated at 37°C for 3 h with 28 U Benzonase and 1:15 (w/w) protease LysC. The concentration of urea was further diluted to 1 M with 50 mM ABC and over-night digestion performed with 1:5 (w/w) trypsin at 37°C. The samples were acidified to a pH of less than 2 with formic acid and desalted with C18 spin cartridges. The cartridges were conditioned with 90% acetonitrile (MeCN), 0.1% formic acid and equilibrated with 5% MeCN, 0.1% formic acid. After loading, samples were washed with 5% MeCN, 0.1% formic acid before elution with 50% MeCN, 0.1% formic acid. All centrifugation steps were performed for 1 min at 1500 g. The eluate was frozen in liquid nitrogen and lyophilized until dry.

Peptides were resolved on a 50 cm C18 chromatographic column on a nLC1000 nano-UPLC and spectra acquired on an Orbitrap Fusion mass spectrometer (both Thermo Scientific). The full chromatographic and mass spectrometry parameters are provided in “MS_parameters” included in Supplementary Data 1. In brief, peptides were resolved by ramping 100% MeCN, 0.1% formic acid against 0.1% formic acid from 3% to 25% in 60 min and further to 32% in 5 min. Precursor scans were acquired in the Orbitrap at 120 000 resolution with an automatic gain control (AGC) target of 2 × 10^5^ on a mass range from 400 to 1500 Th. Fragment scans were obtained for precursors of an intensity of at least 5 × 10^3^ and charge states 2-7 with dynamic exclusion. Higher energy collisional dissociation or electron-transfer higher energy collisional dissociation was used dependent on precursor charge and m/z. MS/MS were acquired in the ion trap at an AGC of 10 000 and fill time of 50 ms.

#### SILAC data analysis

Mass spectrometry data analysis was performed using MaxQuant (v.1.6.1.0) (Cox and Mann, 2008). Parameters specified are provided in “mqpar.xml” included in Supplementary Data 1. The “proteinGroups.txt” output file was post-processed using R to generate “silac_results.csv” (see Supplementary Data 1). Contaminants and decoy-entries were removed and proteins filtered to containing at least two unique peptides. Statistics were based on normalized-ratio entries. For proteins where a SILAC-ratio and the intensity of exactly one SILAC channel was missing, the ratio was imputed by division of the existing intensity by the global minimum intensity. These ratios were normalized through dividing by the median of SILAC ratios of all proteins in the corresponding sample. Finally, a two-sided t-test against a null-hypothesis of mean equal to zero was performed for log_2_-transformed SILAC ratios of *wss1Δddi1Δ+ddi1-D220A* vs. *wss1Δddi1Δ+DDI1-WT*.

Volcano plots from Figures 4B and 5A were generated with R Studio; “log2_fc” values were plotted on x-axis and -log_10_(“pVal”) on y-axis. Lists of 26S proteasomal genes (“Complex: 26S Proteasome complex”, ComplexAc: CXP-2262) and gene ontology “DNA binding” (GO:0003677) were obtained from the Saccharomyces Genome Database (SGD).

To inspect the consistency of quantification of peptides originating from Rpo21 (Figure S5A), an independent quantification approach was used. A peptide library was built from the data-dependent mass spectrometry results in Skyline-daily 4.1.1.18179 (cutoff score: 0.95). In the peptide settings Label:13C(6)15N(2) (K) and Label:13C(6)15N(4) (R) were allowed as isotope modifications. Precursors of library peptides of charges 2-4 were filtered in the transition settings. Three isotopic peaks were extracted within 5 min of their MS/MS identifications. After import of the .raw files the idotp-scores for both the unlabeled and isotopically labelled precursor of Rpo21 (P04050) peptides were generally at least 0.95. The light-heavy-ratios were exported to a .csv file and used to the plots in Figure S5A using R after normalizing to the median label-ratio based on peptides from all proteins.

#### Chromatin fractionation (small scale)

Small scale chromatin extraction was performed from 50 OD of yeast culture growing in log phase using the chromatin extraction protocol described above with the following modifications: cell wall digestion was performed in 1ml of pre-and spheroplasting buffers, the step with 7.5% Ficoll–Sorbitol cushion buffer was skipped. Spheroplasts were lysed in 1.4 ml 18% Ficoll buffer (instead of 14 ml) and no mechanical homogenization was performed. Chromatin pellets were directly resuspended in sample buffer after the EBX wash step. The quality of obtained fractions was controlled by immunoblotting with antibodies against cytoplasmic PGK1 and chromatin H3 proteins.

#### Flow cytometry analysis (FACS)

Cells for FACS analysis (1 ml of cell culture at OD_600_=0.5 or less) were fixed in 70% EtOH and stored at 4°C for up to 1-2 weeks. Fixed cells were centrifuged at 3800 g for 2 min, pellets washed with 300 μl of 50 mM NaCi, pH 7.2, centrifuged for 10 min at 3800 g, resuspended in 250 μl NaCi + 5 μl RNAseA and incubated for 1-3 h at 37°C. DNA was stained with 25 μg/ml propidium iodide for 1 h at 37°C and sonicated 5 times for 5 s with Bioruptor Tween. Flow cytometric analysis was performed in NaCi buffer using a Gallios flow cytometer. FACS profiles were analyzed by Kaluza software.

### Protein purification from insect cells and *in vitro* cleavage assay

Recombinant Ddi1 constructs were produced in insect cells as described in (Spadaro et al., 2017) and summarized below.

#### Cloning of *GST-DDI1* into Bacmid vectors

In-frame GST-tagged *DDI1-WT* and *ddi1-D220A*, constructs were amplified from pFS4148 and pFS4152 respectively using oligos OFS4514 and OFS4515 (see Supplementary Table 1). pFastBac HT A plasmid suitable for the Baculovirus system was modified in order to remove a 6xHis-tag (PCR around plasmid with OFS4512 and OFS4513) and assembled with a respective *DDI1* insert using the NEBuilder^®^ HiFi DNA Assembly Cloning Kit. Assembled products were transformed into DH5α E. coli strains and positive constructs were verified by sequencing. 20ng of each plasmid diluted in water were transformed into DH10Bac bacteria according to manufacturer’s instructions and plated on LB plates containing Kanamycin (50 μg/ml), gentamicin (10 μg/ml), tetracycline (10 μg/ml), X-gal (0.01%) and IPTG (40 μg/ml). Cells were grown for 24h at 37°C and kept at 4°C for 24h to intensify the blue color. A white colony was selected for each construct, grown in LB supplemented with antibiotics and plasmid DNA was purified according to the manufacturer’s protocol. Recombinant Bacmid was further verified by PCR using M13 forward and reverse primers.

#### Transfection of insect cells

2 μg of recombinant Bacmid plasmids were transfected into SF9 insect cells using *Trans*IT^®^-Insect Transfection Reagent (according to the manufacturer’s instructions). A first batch of virus P1 was collected after 72 h and used to re**-**infect P10 plates for 72 h (batch P2). Expression and solubility of recombinant proteins were checked in cells collected from P10 plates. For large-scale expression, insect cells were grown in liquid TC-100 medium supplemented with CD lipid concentrate and infected with the P2 batch of virus for 72 h (5 ml of virus/100 ml of culture).

#### Protein purification

Cells were pelleted by centrifugation at 500 g for 5 min and washed twice with PBS. Pellets were resuspended in 3 ml of LTB buffer (150 mM NaCl, 20 mM Tris-HCl pH7.5, 5 mM EDTA, 1% Triton X-100, protease inhibitor tablet) and incubated on ice for 10 min. Cells were sonicated 4 times for 20 s (30% intensity, Dranson SLPe sonicator) and kept further on ice for 10 min. Lysates were centrifuged for 10 min at 18000 g and soluble fractions containing recombinant proteins were collected and incubated for 2 h at 4°C with 250 μl of Glutathione Sepharose packed beads pre-washed with PBS and equilibrated in LTB buffer. Beads containing bound proteins were washed 3 times with PBS and proteins were eluted 3 times with 250 μl of buffer containing 20 mM reduced glutathione. Eluted proteins were dialyzed for 24 h at 4°C against Prescission Cleavage buffer (50 mM Tris HCl, pH 7.5, 100 mM NaCl, 1 mM EDTA, 1 mM DTT) and cleaved with Prescission protease (10 U/eluate) overnight at 4°C. Prescission protease was removed from the eluate by incubation with 100 μl of packed Glutathione Sepharose beads and the unbound fraction containing recombinant protein was collected. Protein concentration and cleavage efficiency were estimated by SDS-PAGE using BSA as a standard.

#### *In vitro* Top1 cleavage assay

A 32 nt dsDNA stock (1 μg/μl) was prepared by mixing 32bp_nt_templ_For and 32bp_nt_templ_Rev in 10 mM Tris pH=7.5, 50 mM NaCl, boiling for 5 min and leaving at RT for 10 min for re-annealing. Recombinant human Top1 protein (300 ng/reaction) was used as a substrate to test enzymatic activity of recombinant Ddi1 constructs purified from insect cells. The reactions were performed in 25 mM Tris-HCl pH=7.5, 150 mM NaCl in a total volume of 15 μl and mixed in the following order: buffer, 0.2 μg 32nt dsDNA, 250 ng of mono-or 1 μg of poly-ubiquitin, 300 ng Top1, 500 ng recombinant Ddi1 and equilibrated for 2 h at 30°C. DNA was digested with 25 U benzonase and 3 mM MgAc for 15 min at 37°C and reactions were stopped with 5 μl of 4x sample buffer with DTT and 10 min boiling. Samples were resolved on 10% SDS-PAGE gel and silver stained.

#### Silver staining

To perform silver staining, SDS-PAGE gels were fixed by two sequential 30 min incubations in 10% TCA, 30 min in methanol:acetic acid:H_2_O = 50:12:38, washed twice for 20 min in water and fixed for 30 min in 50mM Na_2_B_4_O_7_ and 1**%** glutaraldehyde. Gels were washed three times for 20 min in water, stained for 20 min with 0.8% AgNO_3_ dissolved in 0.5% NH_4_OH and 0.0184 N NaOH and washed twice in water for 7 min. Staining was revealed by 5-10 min incubation in reducing solution (10% EtOH, 0.006% citric acid, 0.0093% formaldehyde); the reaction was stopped with 1% citric acid. Gels were washed with water and dried prior to imaging.

### Chromatin immunoprecipitation (ChIP)

For ChIP with no crosslink, yeast cultures were pelleted, washed twice with cold 1x PBS and directly frozen in liquid nitrogen. To monitor Ddi1 recruitment to the *FRT* locus, cultures were fixed for 15 min with 1% formaldehyde, neutralized by 250 mM glycine for 5 min at room temperature and 10 min on ice, washed twice with cold 1x PBS and frozen in liquid nitrogen. Cell pellets were resuspended in FA buffer (50 mM HEPES-KOH, pH 7.5, 140 mM NaCl, 1 mM EDTA, 1% Triton X-100, 0.1% sodium deoxycholate (w/v)) supplemented with freshly added protease inhibitor. Lysates were mixed with glass beads and homogenized at 4°C in MagNA Lyser 4 times for 30 s at 6000 rpm with 1 min pause. Recovered pellets were centrifuged for 30 min at 18000 g at 4°C. The soluble fraction was discarded and pellets were resuspended in 1 ml fresh FA+inhibitor buffer. DNA was sonicated 20 cycles of 30s-30s in Bioruptor Twin and insoluble fractions were pelleted by 15 min centrifugation at 18000 g at 4°C. Protein concentration was measured by Bio-Rad protein assay and equivalent quantities of total protein (typically 0.5-1 mg) were used for chromatin immunoprecipitation. 1/10 (input) was kept separately at the same conditions as ChIP samples. Lysates were incubated with 1 μl of antibodies against HA epitope tag (to bind Flp-H305L-3HA) or Dynabeads™ Pan Mouse IgG (to directly bind Ddi1-TAP via Protein A) overnight at 4°C with rotation. Antibodies were then coupled to protein G Dynabeads for 3 h at 4°C. Washes were performed using a magnetic separator with 500 μl of FA buffer, twice with the same volume of FA-500 (50 mM HEPES-KOH, pH 7.5, 500 mM NaCl, 1 mM EDTA, 1% Triton X-100, 0.1% sodium deoxycholate (w/v)), twice with buffer III (10 mM Tris-HCl pH 8, 1 mM EDTA, 250 mM LiCl, 1% IGEPAL, 1% sodium deoxycholate (w/v)) and once with TE. Complexes were eluded from beads by two sequential incubations with 100 μl elution buffer B (50 mM Tris-HCl pH 7.5, 1% SDS, 10 mM EDTA) at 65°C. ChIP and input samples were incubated for 2 h at 42°C with 0.75 mg/ml Proteinase K and decrosslinked for at least 12 h at 65°C. DNA was mixed with 1 ml NTB from the Nucleospin Gel and PCR clean up kit and purified according to manufacturer’s recommendations. DNA was eluted from the column twice with 30 μl sterile H_2_O. qPCR reactions were performed using SYBR Select Master Mix with oligonucleotide pairs listed in the key resource table.

### Functional analysis of 26S proteasome activity using fluorescent reporters

GFP fluorescence proteasome assays are based on (Heessen et al., 2003). Plasmids carrying Ub-R-GFP, Ub-M-GFP or G76V-GFP reporters or pYES2 empty vector were co-transformed with pAD54-based Ddi1 constructs or pAD54 empty vector. SC-ura-leu medium was used to retain select for pYES2 and pAD54 plasmids. SC medium was supplemented with 2% galactose to induce expression of GFP reporters. Non-induced cultures were grown in 2% glucose. Live cells were subjected to flow cytometric analysis (FACS) using the FL1 channel of a Gallios flow cytometer. X-mean values used for data analysis were obtained with the Kaluza software. For each genotype X-mean FL1 intensity of the strain carrying empty vector or the reporter was calculated. The empty vector values were subtracted from the intensities of the reporters. Adjusted values were further normalized to the Ub-M-GFP intensity (a control reporter which does not carry a degradation signal) and plotted using Prism 8.

### Measurement of 26S proteasome activity (Suc-LLVY-AMC hydrolysis)

The Suc-Leu-Leu-Val-Tyr-AMC (Suc-LLVY-AMC) fluorogenic substrate for 20S proteasome was used to evaluate chymotrypsin-like activity of 26S proteasome in total cell extracts and in the chromatin fraction. Chromatin was isolated as described in the SILAC MS section. 30 μl WCE aliquots were mixed with 300 μl of EBX without inhibitors to evaluate proteasome activity in total cell extracts. Protein concentration was measured by Bio-Rad protein assay and all samples were adjusted to the same concentration. Final chromatin pellets were washed in 500μl Buffer P (50 mM Tris-HCl pH 7.5, 1 mM EDTA, 1 mg/mL BSA, 1 mM ATP, 1 mM DTT). 25-50 μg of total protein in Buffer P were supplemented with 1 mM ATP and 12.5 μM of Suc-LLVY-AMC substrate. 20 μM MG132 was added to control samples to evaluate basal proteolysis activity. Fluorescence kinetics were measured for 2 h at 37°C with 5 min intervals using Cytation 3 Cell Imaging Multi-Mode Reader (BioTek). Values from the basal activity (+MG132 samples) were subtracted to yield relative fluorescence units.

### Rpb1 stability experiment

Rpb1 degradation after UV treatment was monitored as in (Verma et al., 2011). Briefly, overnight cell cultures were diluted and grown in liquid YEPD medium to an OD_600_=1.0. Cells were then collected and resuspended in 80% of the initial culture volume in the same medium. Liquid cultures (60ml) were then exposed to a UV dose of 400 J/m^2^ in petri dishes in a UV Stratalinker 2400. Cells were either collected directly after treatment or incubated at 30°C in the dark to monitor the recovery in medium containing 100 μg/ml. of cycloheximide (CHX) Cell harvesting was followed by protein extraction and immunoblotting with anti-Rpb1 or anti-PGK1 (internal control). Fluorescent secondary antibodies were used to quantify of relative Rpb1 levels.

Rpb1 turnover analysis in hydroxyurea (HU) was carried out similarly to UV degradation, except that cells were grown to an OD_600_=0.8 before treatment, and HU was directly added to the medium to a final concentration of 200 mM, along with 100 μg/ml of cycloheximide.

## QUANTIFICATION AND STATISTICAL ANALYSIS

Transposon screen data analysis was performed as described in (Michel et al., 2017). The snapshot from the genome browser was made using the generated .wig files uploaded to the UCSC genome browser (http://genome.ucsc.edu/) (Kent et al., 2002). Volcano plots from the Figure 1 were generated using R Studio software. The log_2_ ratio between *tdp1wss1+ auxin* and a pool of other libraries published in Michel et al. 2017 (x-axis), and respective p-values calculated for each yeast gene (y-axis) are plotted. Gene body (Figure 1C) is defined as the ORF. Promoter (Figure 1D) is defined as a 200-bp interval upstream of the ATG translation start site. Analysis of proteomic data is detailed in the “SILAC mass spectrometry” section of STAR methods. To generate Figures 4C and S4C normality was first tested for non-logarithmic and log_2_ values by D’Agostino & Pearson test using Prism 8. Since most of groups did not pass the test for normal distribution, data were further analysed by Mann-Whitney test. Statistical details of other experiments can be found in figure legends. Prism 8 was used to quantify p-values.

## DATA AND SOFTWARE AVAILABILITY

The SATAY sequencing data have been deposited in the European Nucleotide Archive (ENA) under ID code PRJEB31382.

## KEY RESOURCES TABLE

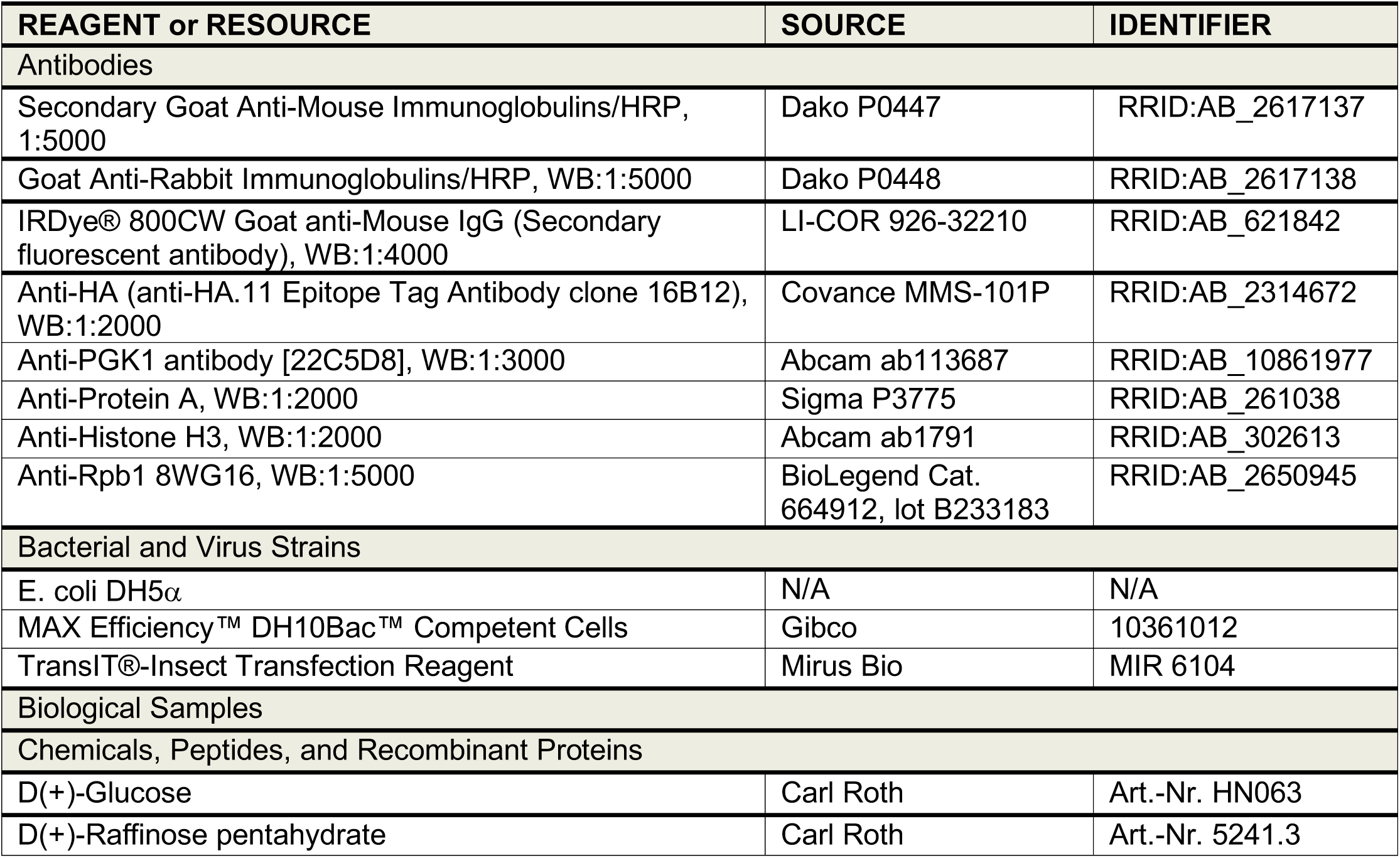
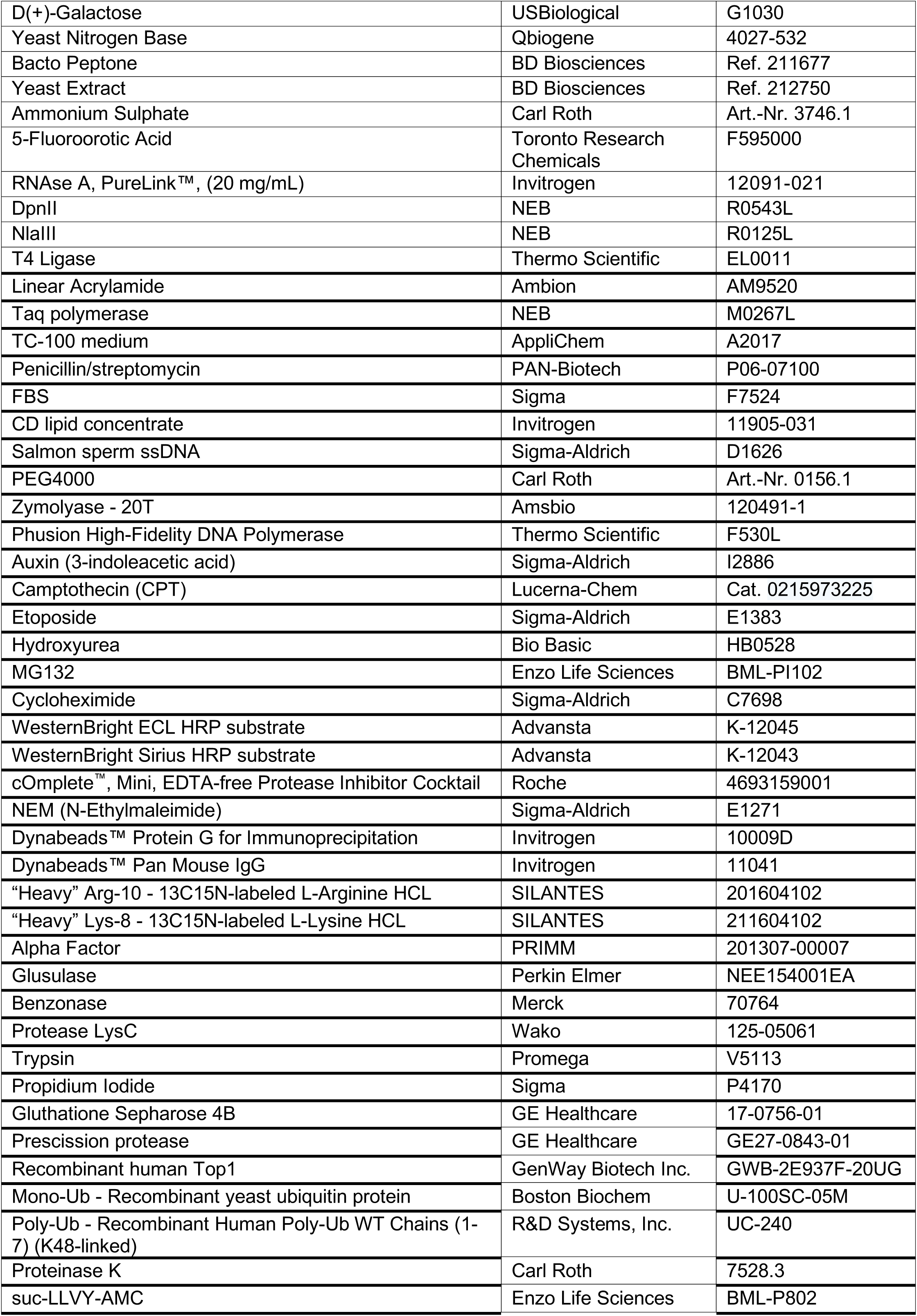
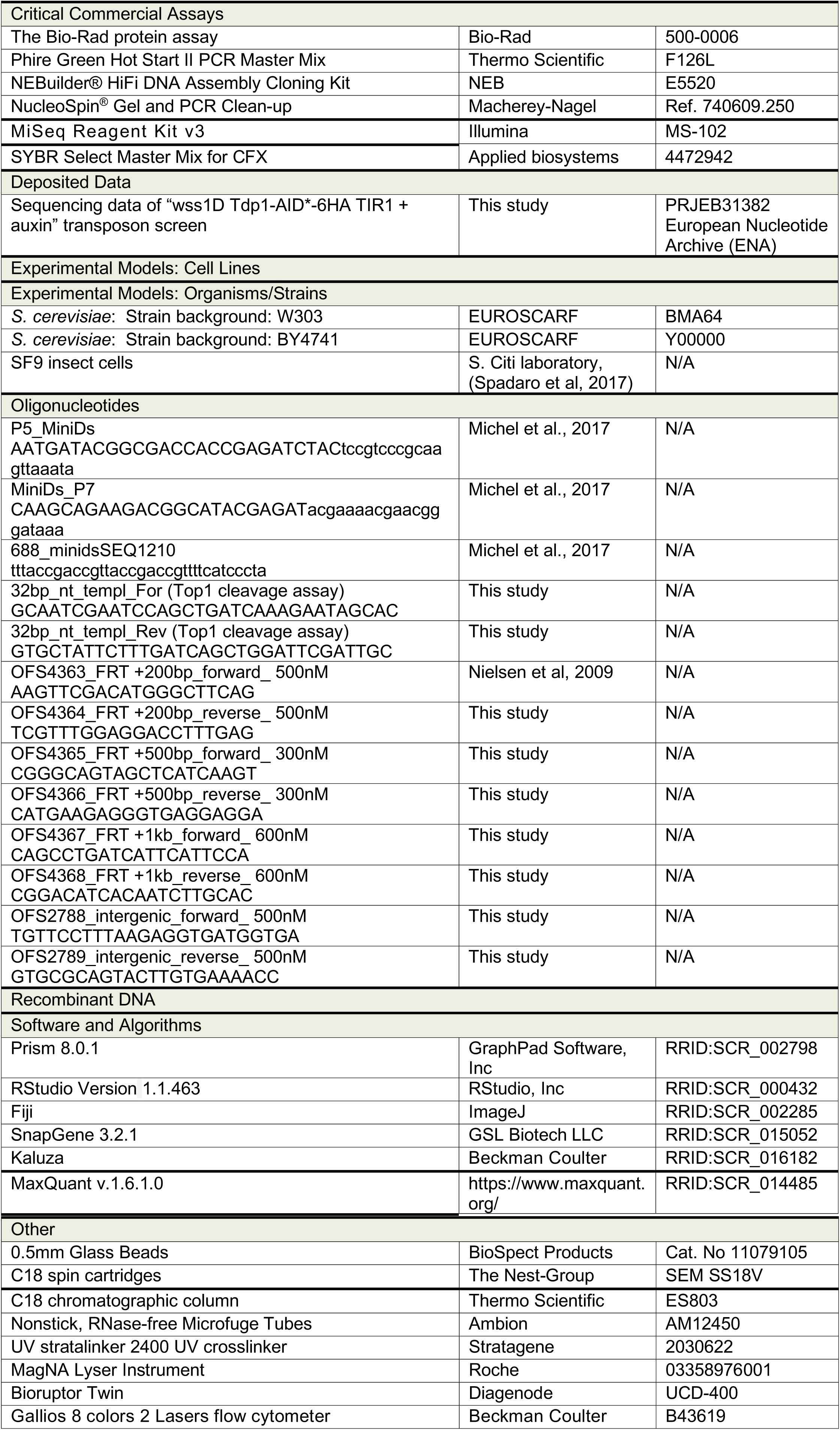

## Supplemental Information

Supplementary Figures 1-5

Supplementary Table 1 – Strains, plasmids and oligonucleotides used in this study Supplementary Data 1 – Data analysis and results of SILAC mass spectrometry related to Figure 4 (MS_parameters.xps, mqpar.xml, silac_results.csv)

## Supplemental Figure Legends

**Supplementary Figure 1:**
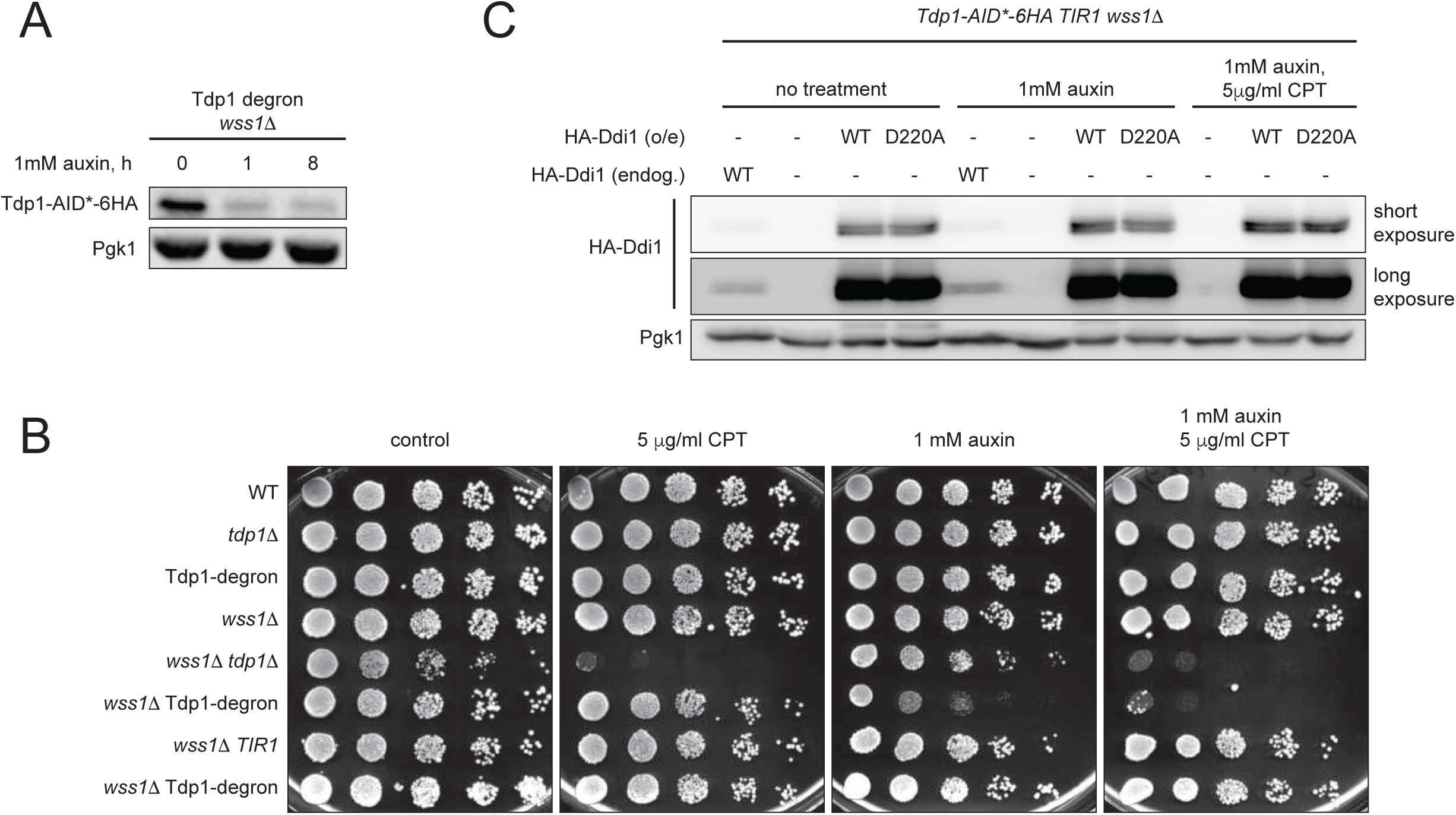
(A) Dynamics of auxin-inducible Tdp1-AID*-6HA degradation in BY4741 genetic background used for transposon screen described in Figure 1B. Efficiency of Tdp1 degradation was revealed by immunoblotting. (B) The Tdp1-degron (*Tdp1-AID*-6HA pADH-TIR1*) system mimics the phenotypes of *tdp1*Δ. 10x serial dilutions of yeast exponential cultures were spotted on YEPD plates supplemented with 1 mM auxin and 5 μg/ml camptothecin (CPT). (C) HA-Ddi1 protein levels in Tdp1-degron *wss1*Δ related to Figure 1G. To allow the comparison with non-treated samples, lanes 1-4 show the same immunoblot membrane as in Figure 1G. Cell overexpressing Ddi1 (o/e) were grown in SC-leu; 1mM auxin was added for 8 h; CPT (5 μg/ml) was added for 1 h prior to collection of total cell extracts.

**Supplementary Figure 2:**
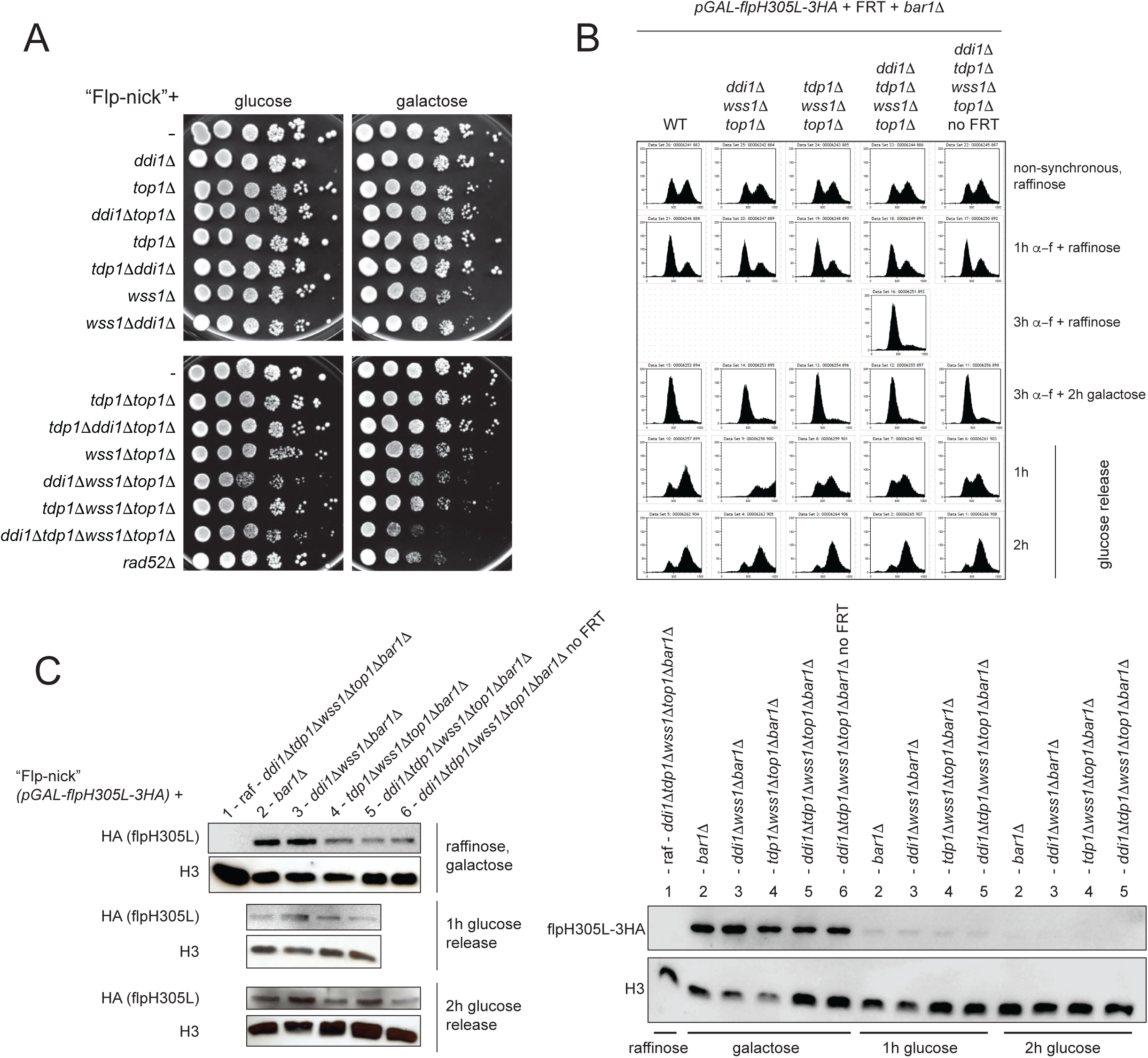
(A) Flp-nick system in different combinations of *ddi1*Δ, *tdp1*Δ, *wss1*Δ and *top1*Δ mutants. The strains were pre-grown in YEP-2% raffinose and spotted on SC-2% glucose (control) or SC-2% galactose medium to evaluate the response to *pGAL-flp-H305L* induction. Plates were imaged after 72 h of growth. (B) FACS analyses of *pGAL-flp-H305L-3HA* strains related to Figure 2E. (C) Total Flp-H305L-3HA protein levels remain similar in different mutant backgrounds, suggesting specific retention of Flp-H305L-3HA at the *FRT* locus. Flp-H305L-3HA protein levels in ChIP input samples were monitored by immunoblotting with anti-HA antibodies. Histone (H3) is used as a loading control. Right panel shows a rapid decrease of total pGAL-flpH305L-3HA protein levels upon transcription repression by glucose.

**Supplementary Figure 3:**
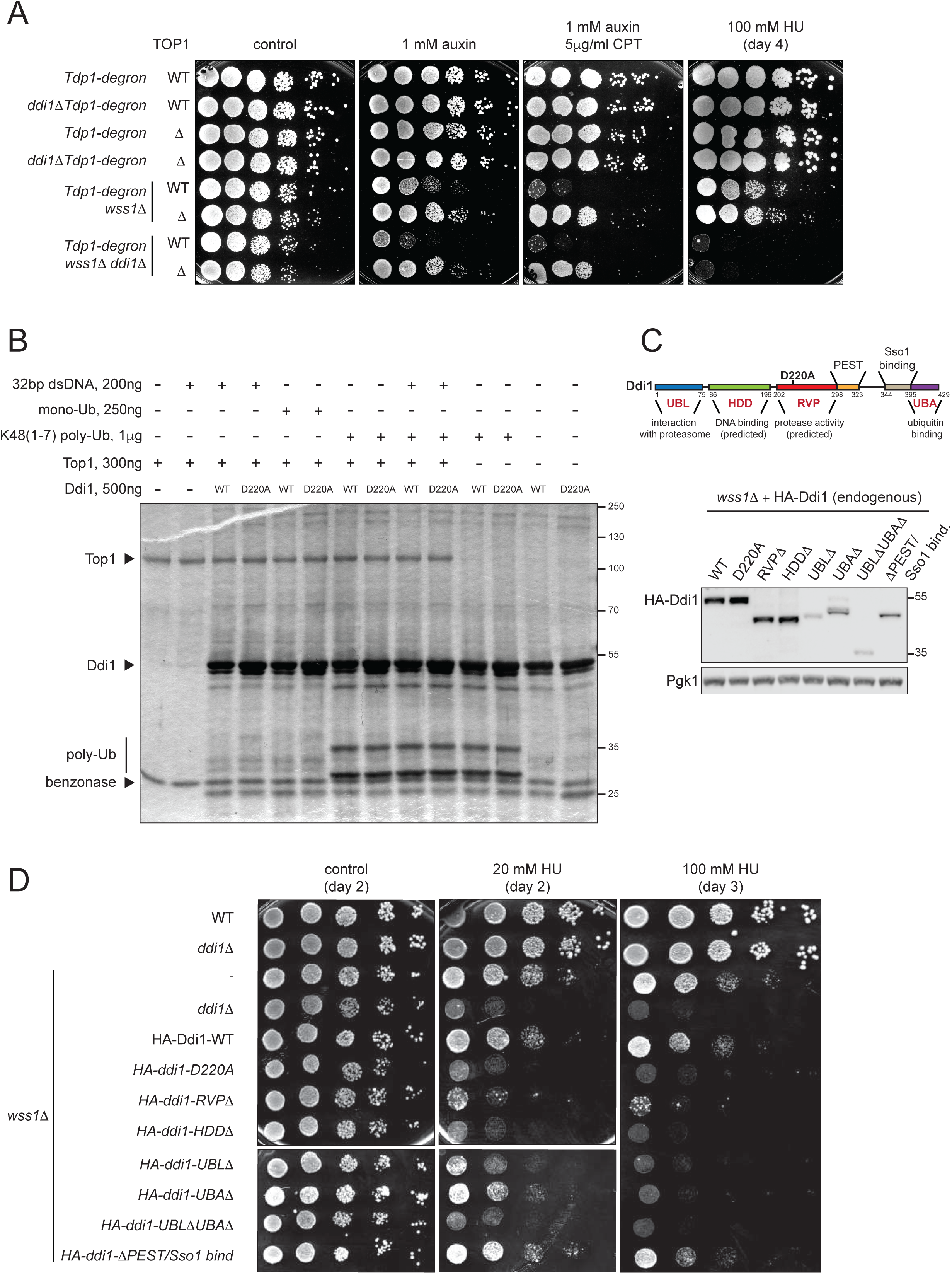
(A) Top1 crosslinks are the main cause of cell death of *tdp1*Δ*wss1*Δ*ddi1*Δ, but not *wss1*Δ*ddi1*Δ mutant in presence of genotoxins. The *top1*Δ specifically improves growth on CPT (Top1 trapping drug), but doesn’t affect HU sensitivity of *wss1*Δ*ddi1*Δ. Tdp1-degron is used to rapidly degrade Tdp1 in presence of auxin (see Figure 1). Cells were spotted on YEPD media supplemented with the indicated drugs; images were taken after 48 h, with the exception of the 100mM HU plate that was scanned on day 4 (96 h). (B) Ddi1 protease activity is not detected *in vitro*. Recombinant Ddi1-WT and Ddi1-D220A were purified from insect cells and used for Top1 cleavage assay (see STAR Methods section for the details). Proteins were resolved by SDS-PAGE and the gel was silver stained. (C) Top: Ddi1 domain organization with additional PEST and Sso1 binding regions. Bottom: UBA and UBL domains affect Ddi1 protein stability. Protein levels of genomic HA-tagged Ddi1 mutants were examined by immunoblotting. (D) HU sensitivity of endogenous *DDI1* mutants shown in (C). Cells were spotted on YEPD medium containing 20 mM or 100 mM hydroxyurea (HU) and imaged after 2-3 days (48-72 h) of growth.

**Supplementary Figure 4:**
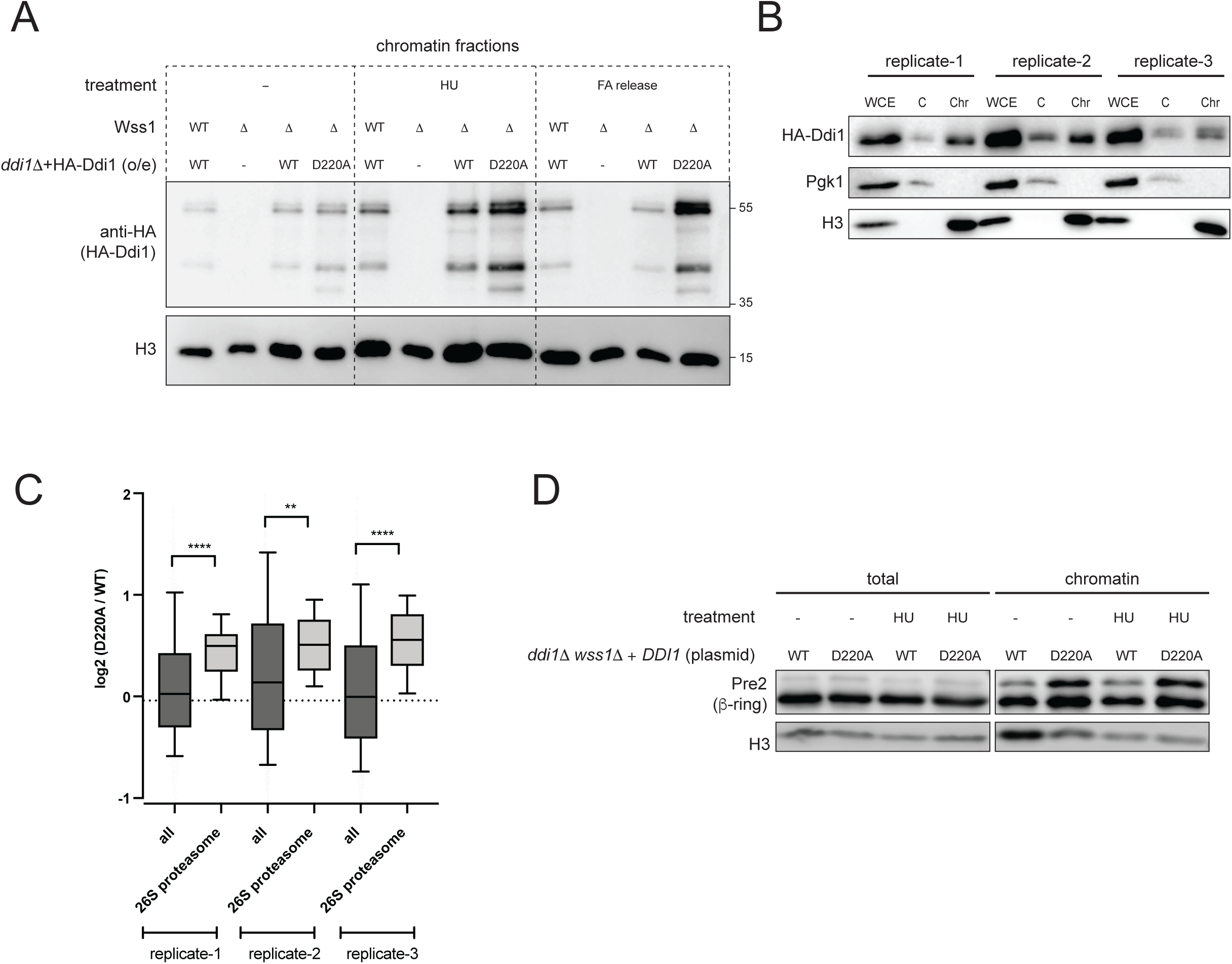
(A) In the absence of Wss1, the catalytic protease mutant Ddi1-D220A accumulates on chromatin after genoxin treatment. Cells were subjected for 2 h hydroxyurea (HU, 200mM) treatment or were released for 2 h from 15min formaldehyde (FA, 40 mM). The protein levels of the overexpressed HA-Ddi1 constructs in chromatin fractions were compared by immunoblot. (B) The quality of chromatin-enriched fractions used for SILAC experiment related to Figure 4A. WCE, whole cell extracts, C, cytoplasm, Chr, chromatin. Pgk1, a marker of a cytoplasmic fraction; H3, a marker of a chromatin fraction. (C) Enrichment of the 26S proteasome complex for each of the three SILAC MS independent replicates described in Figure 4A. The analysis was performed as described in Figure 4C. (D) Validation of 26S proteasome enrichment related to Figure 4D. The 26S proteasome Pre2 subunit is enriched on chromatin in *wss1*Δ*ddi1*Δ in the absence of HU. The experiment was performed as described in Figure 4D.

**Supplementary Figure 5:**
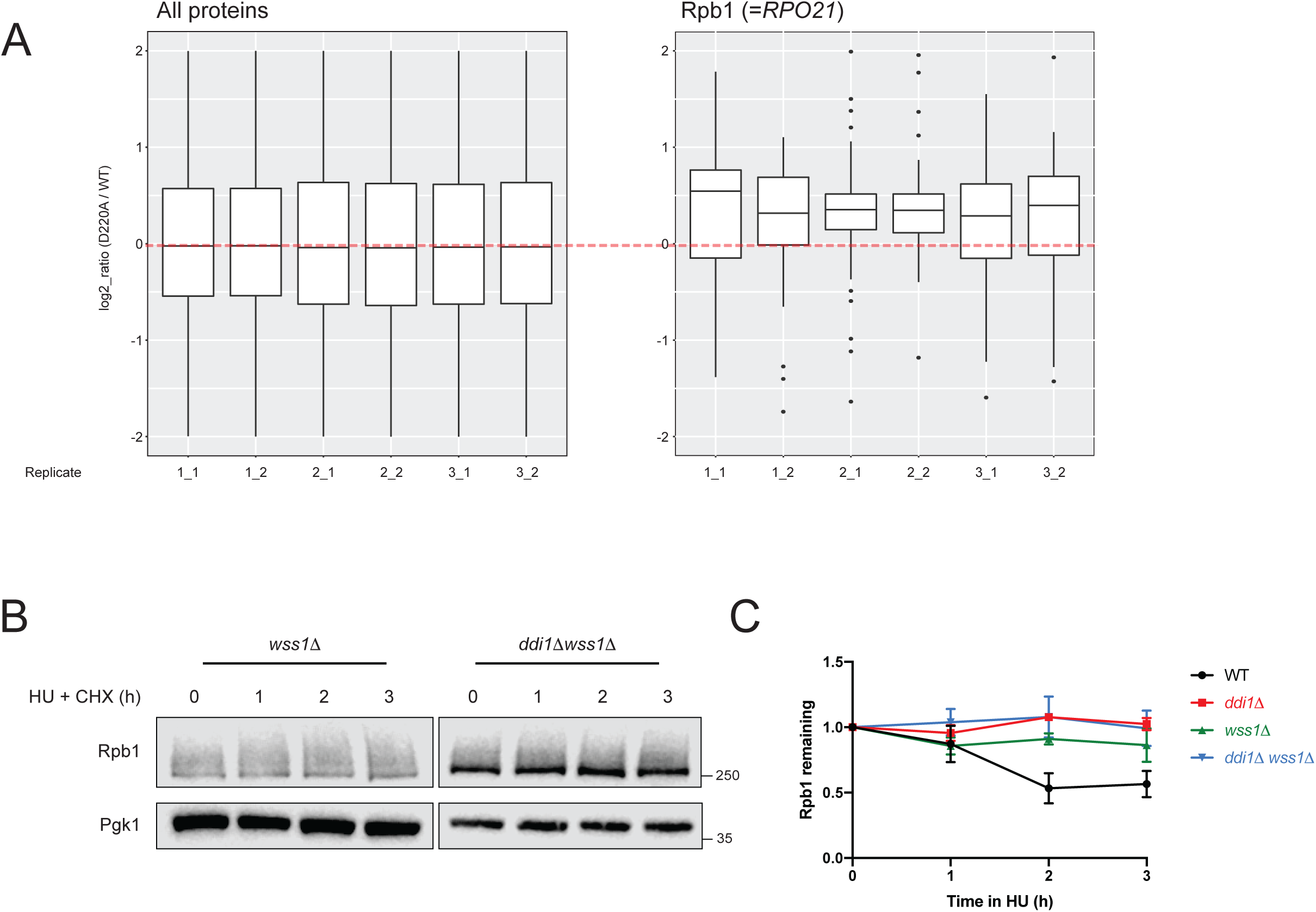
(A) Rpb1 is significantly enriched on chromatin in *wss1*Δ*ddi1*Δ *+ddi1-D220A* mutant. SILAC mass spectrometry data from Figure 4 were analyzed and depicted as boxplots of normalized log_2_ (D220A/WT) ratios for all peptides of 2199 identified proteins (left) and for all identified Rpo21 peptides (right). Boxes represent 25^th^, 50^th^ and 75^th^ percentile. Whiskers extend from hinges to most extreme value no further than 1.5 interquartile ranges. Three biological replicates with two repeat injections are shown on the x-axis. (B-C) Rpb1 degradation is defective in *wss1*Δ mutant. Relative Rpb1 levels were analyzed and quantified as in Figure 5C. The data from Figure 5C (WT and *ddi1*Δ) are also plotted.

